# Integrative analysis reveals generalizable human neurodegenerative disease-associated glial states

**DOI:** 10.1101/2025.09.25.678630

**Authors:** Liam Horan-Portelance, Dominic J. Acri, Alexandra Mann, Janet Brooks, Hannah M. Bailey, J. Raphael Gibbs, Alex R. DeCasien, Mark R. Cookson

## Abstract

Glial cells are known to respond transcriptionally in multiple neurodegenerative diseases (NDDs). In particular, microglial states have been characterized in Alzheimer’s disease and mouse models of amyloidosis as disease-associated microglia. Although single-cell transcriptomic technologies have increased the dimensionality of information available across cell states, few studies have systematically tested for changes in glial transcription across brain regions and disease states. Here, we report a statistical framework for glial annotation, disease association, and transcriptional profiling, which facilitate identification of generalizable glial states that are present across a spectrum of NDDs (Alzheimer’s disease, Parkinson’s disease, amyotrophic lateral sclerosis, and frontotemporal dementia) by re-analyzing data available in four multi-region atlases. We identify seven astrocyte substates, 14 microglia/myeloid substates, and five oligodendrocyte substates where transcriptional variability is attributable to region, disease, or study-specific effects. Regional heterogeneity of astrocytes masked disease associations, even within cortical subregions. We found only limited oligodendrocyte transcriptional heterogeneity, resulting in few substates for further interrogation. Notably, microglia showed the strongest evidence for disease association. We show, for the first time, that this association exists across different NDDs. Using latent factor analysis, we created a consensus human neurodegenerative disease-associated microglia (hnDAM) signature, which we experimentally validated in 11 independent sample series. We demonstrate that the hnDAM signature is a statistically testable biomarker for conserved microglial activation in NDDs by: i) comparing to murine DAM-like signatures, ii) performing transcription factor analysis, and iii) modeling transcriptional reprogramming perturbations in iPSC-derived microglia. Importantly, we find for the first time a way to make direct comparisons between DAM-like activation profiles in separate studies and propose a novel modeling paradigm via PIKfyve inhibition. Taken together, this work broadens our understanding of glial activation across neuropathologies and reveals hnDAM as a putative therapeutic target that can be utilized in any transcriptomic study of patients suffering from NDDs.

**Graphical Abstract:** 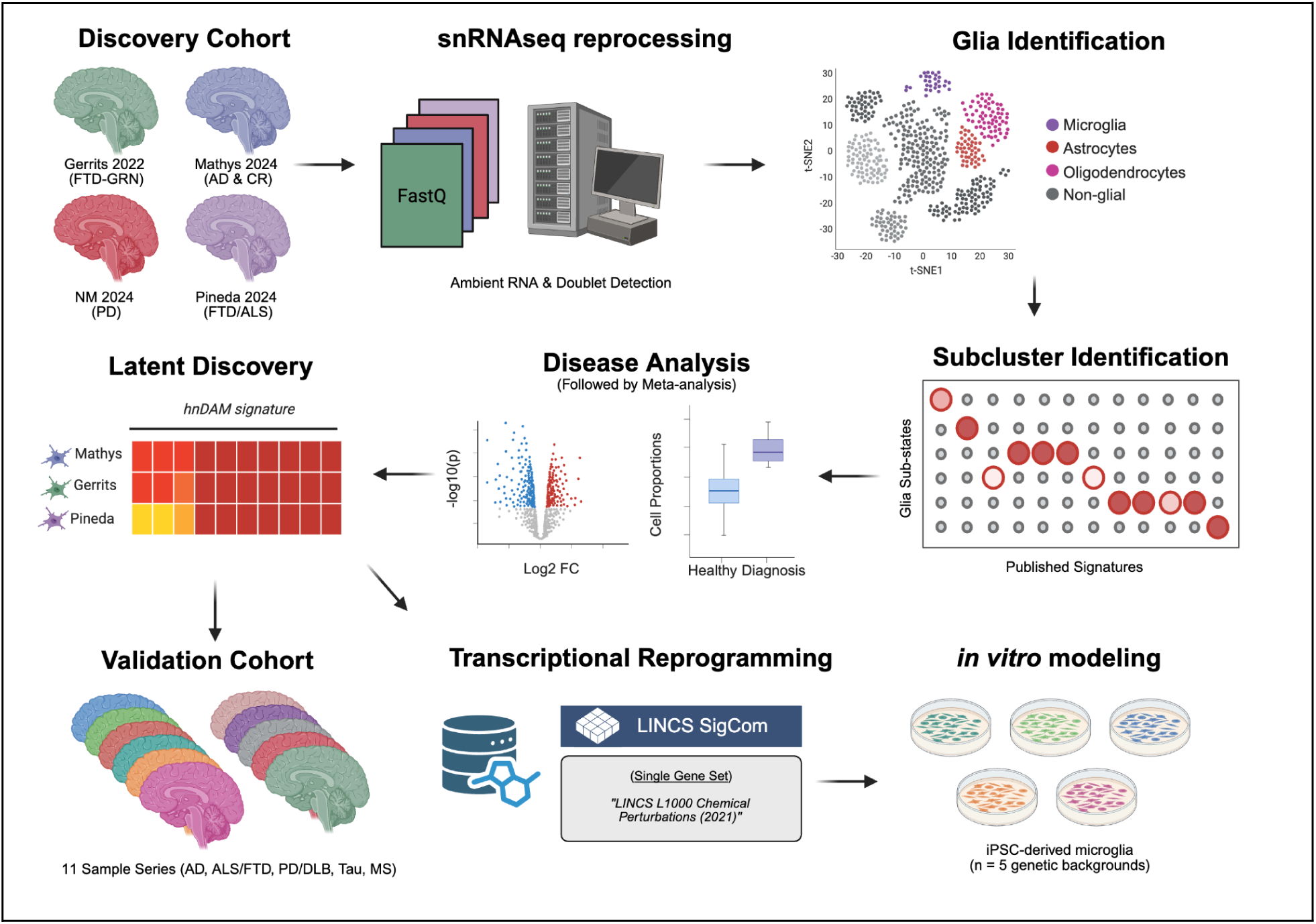

## Introduction

Age-related neurodegenerative diseases (NDDs) are characterized by neuronal loss and protein deposition^1^. Although types of pathology and affected neurons vary across diseases, one unifying phenomenon is the appearance of reactive gliosis^2^. While the functions of glia in the brain are well characterized^3^, defining and replicably describing disease-associated glial subtypes remains a challenge. Advancements in the field of single-cell/nucleus RNA sequencing (sc/snRNA-seq) have allowed deep cell type profiling, but variable naming conventions and strategies to define disease association^4^ have limited replicability across studies.

The major glial cell types present in the human brain are astrocytes, microglia, and oligodendrocytes^3^, and each displays evidence of transitioning between homeostatic, disease-associated, and other activated states in the presence of molecular insults. Microglia present the most evidence of disease-associated transcriptional states. The previous paradigm of M1 (“pro-inflammatory”) and M2 (“anti-inflammatory”) activation states^5^ has been further refined using scRNA-seq technologies to include a spectrum of disease-associated states^4^. Astrocytes have also been described by A1/A2 states^6^, however description of disease-associated astrocyte subtypes in the human brain is largely lacking. Finally, oligodendrocyte responses to pathology have been reported in mice^7–9^, but there is little evidence of similar changes occurring in human postmortem tissue.

A challenge facing glial subtype interrogation is the lack of an agreed-upon taxonomy, as has been developed for neuronal subtypes^10,11^, resulting in study-specific annotations. Many studies have relied on one or a small handful of marker genes discovered in mouse single-cell studies to claim presence or absence of human disease-associated cell types. Given that claims made on single sample series of single-cell data may be limited by sparsity and statistical power^12^, there is an urgent need for robust discovery integrating independent sample series. Several recent studies have either leveraged meta-analysis^13,14^ or primary analysis of newly-collected sample series comprising multiple NDD subtypes^15–17^ to describe aspects of disease-associated signatures. However, these studies predominantly profile neocortical Alzheimer’s disease tissue, so there remains a need for disease association testing in a unified manner across the spectrum of NDDs.

Because there is still a gap in the literature for defining NDD-associated glial subtypes, we aimed to create a unified cohort of cross-NDD affected glia. To this end, we integrated each glial cell population from four multi-region atlases of NDDs^18–21^. Although larger donor sampling cohorts have been recently published, we aimed to include multiple sample series from matched donors to specifically power discovery across regions. Critically, all data were re-processed together with ambient RNA correction, doublet filtering, and batch correction. We found the largest regional heterogeneity in astrocytes and the strongest disease-associated effects in microglia. Following meta-analysis of cell subtype proportions and differential expression in all glial types, we report predominantly shared, but several disease-specific effects. To further characterize pan-NDD microglia effects, we performed latent factor analysis to define a human neurodegenerative disease-associated microglia (hnDAM) signature that we then tested in a replication cohort of 11 independent sample series. Furthermore, we performed *in silico* transcriptional reprogramming followed by *in vitro* modeling in iPSC-derived microglia to experimentally recapitulate hnDAM signatures. We show that the hnDAM signature can be mimicked by exposure to neuronal debris, along with HDAC, PIKfyve, or BTK inhibition. Taken together, our data provides evidence for pan-NDD effects in glia and demonstrates related aspects of microglial activation *in vitro*.

## Results

### Glial transcriptional diversity

To investigate the transcriptional diversity of three major glial cell types, we re-processed snRNA-seq data from four studies spanning NDD diagnosis, brain region, and sex^18–21^ (**Table S1;** abbreviation keys in table). We implemented a rigorous quality control pipeline to correct for ambient RNA, detect and remove doublets, and annotate glial signatures (**Fig. S1A-D**, **Table S2-5**). Clustering revealed seven astrocytic, 14 microglial, and five oligodendrocytic subtypes (**Fig. 1A**). Cell counts from subclusters fit a negative binomial distribution (**Table S6;** mean = 9,831 ± 593, size = 0.26 ± 0.02) and were determined by sketch-based integration to preserve rare cell types^22,23^.

**Fig. 1.**
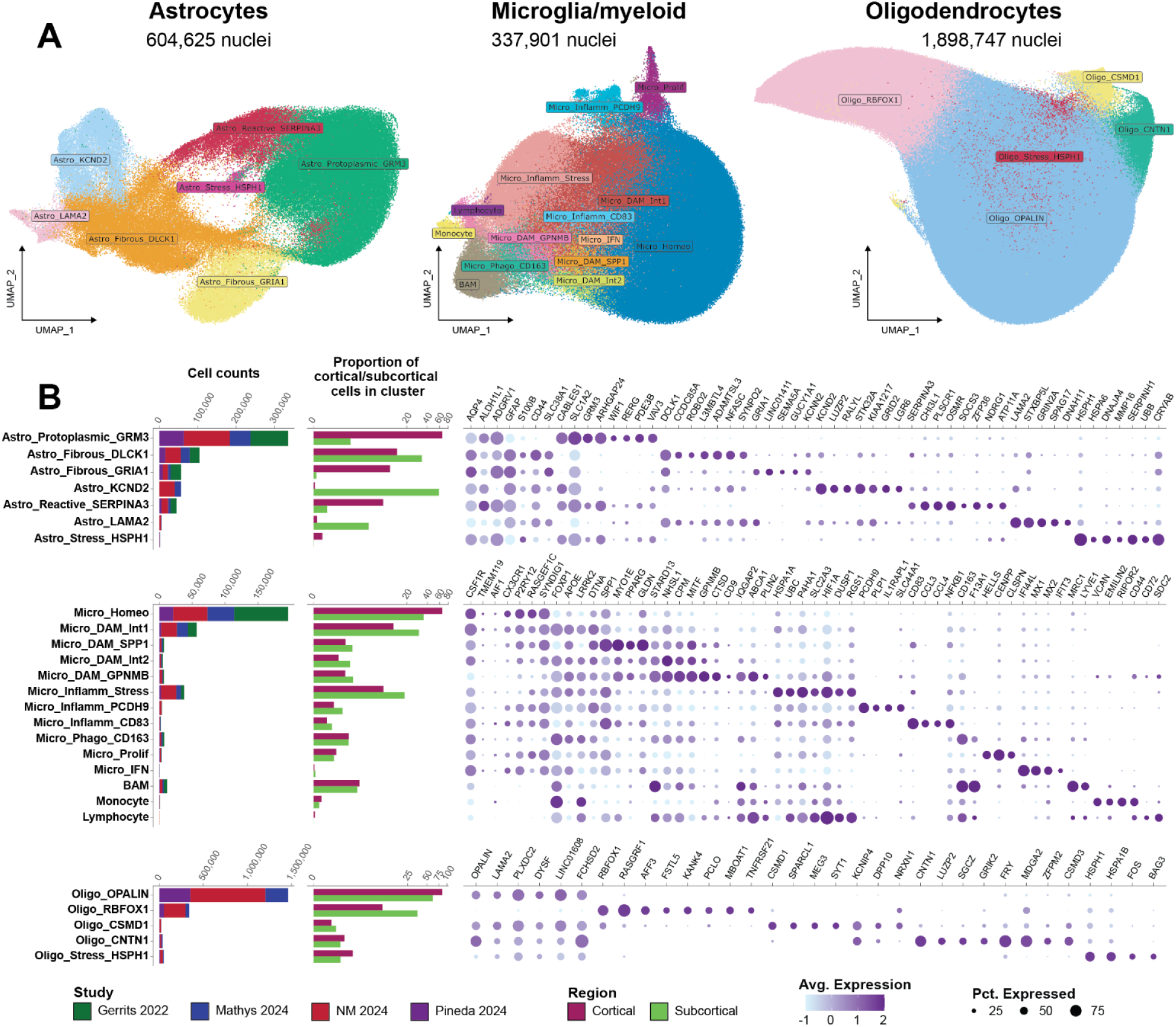
Overview of glial subtype and regional diversity. (**A**) UMAPs of glial cells from atlas-scale NDD datasets^18–21^ annotated using previously published human and mouse substates (**Fig. S2**). (**B**) Study-level statistics and cell type identity metrics report the cells recovered, per-cluster cell assignment to cortical or subcortical regions, and expression of subcluster-specific marker genes.

To confirm the identity and source of glial subtypes, we quantified cell count per condition, proportion of cells from cortical or subcortical brain regions belonging to each cluster, and specific marker genes (**Fig 1B**, **Table S6-9**). Protoplasmic astrocytes (*GFAP*^Low^)^24^, homeostatic microglia (*P2RY12*^High^, absence of activation markers), and *OPALIN*+ oligodendrocytes made up the majority of human brain glial cells. This was followed by fibrous (*GFAP*^High^) astrocytes, an intermediate DAM-like microglial subtype (Micro_DAM_Int1: *APOE*^High^/*SPP1*^Low^/*GPNMB*^Low^), and *RBFOX1*+/*RASGRF1*+ oligodendrocyte populations for each respective glial type. Notably, astrocyte subclusters were highly transcriptionally diverse across brain regions, with four of the seven reporting significant proportion differences between cortical and subcortical regions (Astro_Fibrous_GRIA1: t(8.00) = 3.41, p = 0.009; Astro_Reactive_SERPINA3: t(8.14) = 2.87, p = 0.020; Astro_Protoplasmic_GRM3: t(8.03) = 2.79, p = 0.024; Astro_KCND2: t(2.00) = -5.739, p = 0.029). To provide further resolution of astrocytes, we subclustered cells derived from cortical and subcortical regions separately, yielding six cortical clusters and nine subcortical clusters (**Table S11**). While cortical astrocytes were modestly separated by neocortical or allocortical identity (**Fig S2A-C**), subcortical astrocyte identity was highly impacted by regional differences between the DMV, GPi, and TH (**Fig S2D-F**). In contrast to astrocytes, we found that no subtypes of microglia or oligodendrocytes were nominally significant for regional effects on cell counts.

### Disease-associated glial subtypes

Next, we evaluated the association of glial subtypes with NDD status. Proportions of each subtype in each sample ([donor x region]) were modeled against disease state with a generalized linear model (GLM) (**Table S12**). We used multivariate adaptive shrinkage to identify patterns of sharing across datasets, refine the beta values from the GLMs, and estimate local false sign rates^25^ (LFSRs) (**Fig. 2A, Table S13; see “Cell subtype proportion analysis”**). Notably, we observed stronger disease association of microglial subtypes compared to cortical astrocyte or oligodendrocyte subtypes (**Fig. 2A**). Specifically, we identified four microglial subpopulations (Micro_DAM_Int1, Micro_DAM_SPP1, Micro_DAM_Int2, and Micro_DAM_GPNMB) which displayed strong disease association across multiple NDDs and brain regions (**Fig. 2B**). Additionally, homeostatic microglia (Micro_Homeo) were significantly depleted across several NDDs (i.e., FTD-GRN and ALS/FTLD) and regions.

**Fig. 2.**
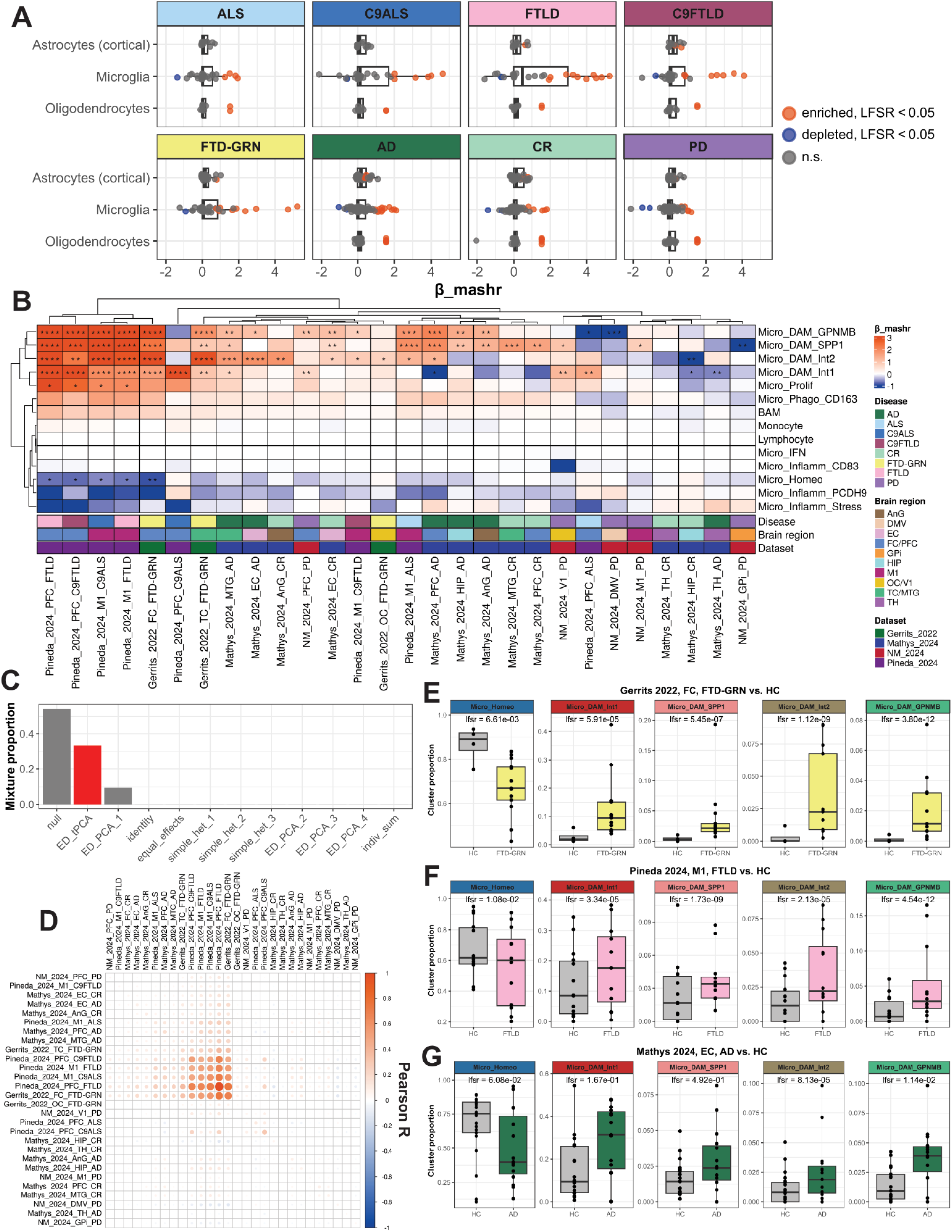
Proportions analysis reveals presence and absence of disease-associated glial subtypes. Differential abundance of subtype populations was tested using a GLM covaried by sex and age. β and LFSR values were refined and estimated using *mashr*^90^. (**A**) Summary of *mashr*-refined β values. Each point represents a comparison between diagnosis and control for a single subtype within a given brain region. Subtypes significantly enriched or depleted across NDD conditions (*LFSR* < 0.05) are colored by effect direction. (**B**) Microglial differential abundance reveals 4 disease-associated subtypes across NDD and regions (**** LFSR *< 0.0001,* *** LFSR *< 0.001,* ** LFSR *< 0.01,* * LFSR *< 0.05*). (**C**) Estimated mixture proportions for different covariance matrices generated from *mashr* analysis of microglia reveals that a data-driven pattern (“ED_tPCA”) explains the most sharing (other than null effects). (**D**) Covariance matrix demonstrates patterns of shared variance explained by ED_tPCA (capturing shared effects across NDDs on the ALS-FTD spectrum). (**E-G**) Per-sample proportion changes of DAM-like subtypes in FTD-GRN, FTLD and AD. See **Table S13** for summary statistics.

We next examined patterns of shared variance in proportion changes across the disease by region comparisons. While most of the variance was explained by null effects, a covariance pattern discovered via extreme deconvolution (“ED_tPCA”) was the next most common pattern in the data (i.e., “ED_tPCA” had the second highest mixture proportion, which is a weight that represents how common a pattern of sharing is across features) (**Fig 2C**). This “ED_tPCA” pattern primarily reflected shared effects across NDDs on the ALS-FTD spectrum (**Fig. 2D**), including conditions from both the Gerrits_2022 (FTD-GRN) and Pineda_2024 (ALS/FTLD) datasets. Importantly, microglial disease associations revealed by *mashr* analysis were also reflected in the donor-level raw proportion changes across studies (**Fig. 2E-G**).

Strikingly, neither cortical astrocyte nor oligodendrocyte subtypes displayed similarly strong nor conserved disease associations compared to microglia (**Fig. S3**). We detected enrichment of two *GFAP*+ cortical astrocyte subpopulations (Astro_Fibrous_CD44 and Astro_Fibrous_GRIA1) across several brain regions and disease subtypes (**Fig. S3A, B**). Additionally, we observed one disease-associated oligodendrocyte subpopulation (Oligo_Stress_HSPH1) (**Fig. S3C**). Notably, this association seems to be driven by outliers at the donor level (**Fig. S3D**), and this population makes up only 2.46% of all recovered oligodendrocytes (46,719 Oligo_Stress_HSPH1 of 1,898,747 total oligodendrocytes). These results indicate that, out of all glial cell types present across these NDD atlases, microglia have the strongest evidence of showing shared responses to disease.

### Shared responses to neurodegenerative diagnoses within cell types

To investigate how glial cells respond in NDDs, we performed differential expression, making the same comparisons done for glial proportions. We employed a differential expression strategy with latent covariates (See **“Differential gene expression analysis”**; **Fig. S4A-B**; **Table S14-16**) comparable to pseudobulk approaches that correct for cell-, donor-, or read-depth artifacts (**Fig. S5A-C**). Notably, the NM_2024 dataset^21^ displayed inflation of differentially expressed genes in pseudobulk, resulting in low spearman correlation coefficients between cell-level and pseudo-bulked analysis in identical comparisons with identical covariates (**Fig. S5D**). Other comparisons showed comparable z-score effect sizes across methods (e.g., Astrocytes_Gerrits_2022_FC_FTD-GRN, rho = 1; **Fig. S5E**). As a result, we proceeded with meta-analysis using cell-level DE results from *nebula*.

We performed differential expression in microglia (**Table S17**) and found 9,730 genes with evidence of responding in at least one comparison (**Table S18;** 18,203 genes tested, LFSR < 0.05). Of these, 8,486 responded in 2 or more conditions and 6,603 responded across conditions in separate studies. We applied multivariate adaptive shrinkage across all conditions in the discovery cohort and found that the most common pattern in the data fit a data-driven covariance matrix (“ED_tPCA”; 70.45% mixture proportion) (**Fig. S6A**). In contrast, condition-specific effects (i.e., patterns of differential expression unique to one comparison only) were much less common (totaling <14.7% mixture proportions). Similar to proportion analyses above, the most common pattern (“ED_tPCA”) reflects shared effects across FTD datasets (**Fig. S6B**). Interestingly, these shared effects of FTD were anti-correlated with the effects of cognitive resilience, indicating that genes best fit to the “ED_tPCA” pattern may explain resilience to dementia in the presence of AD-related neuropathology. When the 2,500 genes whose cross-condition variance is best explained by the “ED_tPCA” pattern were k-means clustered (**Fig. S6C, Table S19**), as opposed to other NDD conditions, clusters one and two were upregulated in cognitive resilience while clusters four and six were downregulated. The set of 981 genes from clusters one and two were enriched for pathways involved in immune activation (**Fig. S6D**; **Table S20**), while the set of 1,085 genes from clusters four and six were nominally significantly enriched for insulin signalling,although these results were not significant after FDR correction (**Fig. S6E**; **Table S20**). These results identify a common response of microglia across NDDs that is negatively correlated to the response in cognitive resilience.

Performing differential expression in astrocytes (**Table S21**), we reported 15,628 genes responsive in at least one comparison (**Table S22;** 21,885 genes tested, LFSR < 0.05). Of these, 14,154 responded in 2 or more conditions and 11,922 responded across conditions in separate studies. Performing meta-analysis across all conditions in the discovery cohort, we found the maximum amount of sharing (48.44% mixture proportion) was explained by a data-driven ED_tPCA covariate model (**Fig. S7A**). In contrast, individual effects made up less than 8.8% mixture proportion. The largest patterns of sharing were across NDDs, with notable anti-correlation between cognitive resilience and FTLD (**Fig. S7B**). Interestingly, PD and ALS shared more similarity with frontotemporal dementia in astrocytes (4 comparisons with corr > 0.3) than those observed in microglia. When the 3,502 genes best explained by the ED_tPCA than any other patterns of sharing were k-means clustered (**Fig. S7C**, **Table S23**), cluster two was uniquely decreased in ALS while cluster five was uniquely increased in cognitive resilience. The set of 312 genes from cluster two was enriched for pathways involved in oxidative phosphorylation (**Fig. S7D**, **Table S24**), while the set of 568 genes from cluster five was enriched for pathways involved in innate immune responses (**Fig. S7E**, **Table S24**). These results show that the patterns of sharing seen in microglia are also seen in astrocytes, including modulation of oxidative phosphorylation. Importantly, there appears to be a subset of shared DEGs that replicate the immune activation in cognitively resilient individuals also seen in microglia.

Performing differential expression in oligodendrocytes (**Table S25**), we reported 15,401 genes with evidence of responding in at least one comparison (**Table S26;** 18,905 genes tested, LFSR < 0.05). Of these, 14,246 responded in 2 or more conditions and 10,371 responded across conditions in separate studies. By performing meta-analysis across all conditions in the discovery cohort, we reported the maximum amount of sharing (55.40% mixture proportion) is explained by a data-driven ED_tPCA covariate model (**Fig. S8A**). In contrast, individual effects made up less than 5.1% mixture proportion. The largest patterns of sharing were between FTD conditions and PD. In contrast, effects within ALS and cognitive resilience contained less sharing with other NDDs (**Fig. S8B**). There was little sharing between ALS comparisons and any non-ALS comparisons in oligodendrocytes from the discovery cohort (0 comparisons with corr > 0.3) in contrast to microglia and astrocytes (18 and 6, respectively). When the 5,533 genes that contributed more to the ED_tPCA than any other patterns of sharing were k-means clustered (**Fig. S8C**, **Table S27**), 1,238 genes of cluster two were uniquely downregulated in ALS. Similar to astrocyte cluster two, oligodendrocyte cluster two genes were enriched for pathways involved in aerobic respiration (**Fig. S8D**, **Table S28**). No k-means clusters were uniquely up-regulated in ALS comparisons, however cluster three is weakly increased in ALS and select regions from cognitively resilient individuals. This cluster was enriched for terms involved in cellular trafficking and morphogenesis (**Fig. S6E**, **Table S28**). Collectively, these results suggest that oligodendrocytes respond similarly across NDDs with the exception of a uniquely downregulated oxidative phosphorylation signature in ALS.

Notably, across all three glial cell types, there was little or no response in the visual cortex of individuals with PD. This is consistent with limited pathology in this region in PD^26,27^, and its inclusion as a negative control in the original study^21^. Importantly, the preservation of null findings in this region across astrocytes, microglia, and oligodendrocytes suggest that our differential expression strategy minimizes false discovery.

### Latent factorization uncovers cross-disease human neurodegenerative disease-associated microglia (hnDAM) signature

Given the existence of four seemingly transcriptionally similar NDD-associated microglial subtypes, we next asked whether there may be an underlying “disease-associated microglia” signature that unifies these subtypes across NDDs. We therefore performed latent factorization on a per-study basis. The FTD-GRN discovery cohort (Gerrits_2022) revealed 16 gene expression programs (GEPs), where GEP5 was specific to Micro_DAM_SPP1, Micro_DAM_Int2, and Micro_DAM_GPNMB (**Fig. S9A**, **Table S29**). The AD discovery cohort (Mathys_2024) revealed 12 GEPs, where GEP3 was specific to Micro_DAM_SPP1, Micro_DAM_Int2, and Micro_DAM_GPNMB (**Fig. S9B**, **Table S30**). The ALS/FTLD discovery cohort (Pineda_2024) revealed 12 GEPs, where GEP5 was specific to Micro_DAM_SPP1, Micro_DAM_Int2, and Micro_DAM_GPNMB (**Fig. S9C**, **Table S31**). The PD discovery cohort (NM_2024) revealed 17 GEPs, where GEP5 was specific to Micro_DAM_GPNMB, Monocytes, and Lymphocytes (**Fig. S9D**, **Table S32**). Based on the stratification of DAM subclusters where either *SPP1* or *GPNMB* are the top markers with the “intermediate 2” DAM subcluster, we selected Gerrits_2022_GEP5, Mathys_2024_GEP3, and Pineda_2024_GEP5 to create a consensus human neurodegenerative disease-associated microglia (hnDAM) signature (**Fig. 3A**). This data suggests that there is an expression program specific to microglial subtypes enriched across NDDs.

**Fig. 3.**
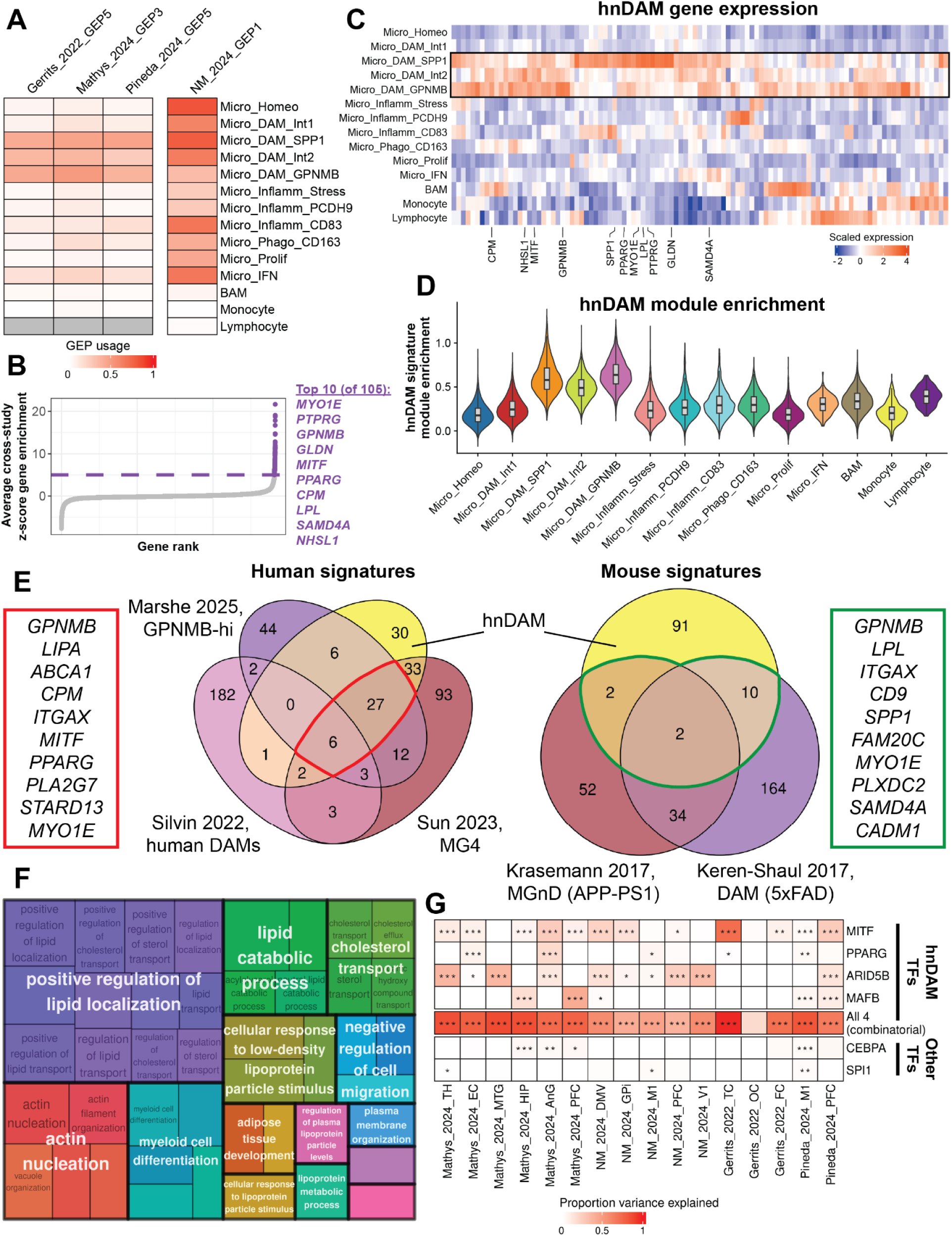
Latent factorization uncovers a 105-gene consensus human neurodegenerative disease-associated microglial (hnDAM) signature. Microglial gene expression programs (GEPs) were defined using *cNMF*^100^ (see “**Microglia signature discovery via latent factorization”**). (**A**) GEPs uniquely enriched in subtypes associated with NDD diagnoses were apparent from three of the four discovery studies. Notably, no GEP from NM_2024 was specific to DAM subtypes (see **Fig. S9D**). (**B**) Genes were prioritized from Gerrits_2022_GEP5, Mathys_2024_GEP3, and Pineda_2024_GEP5 by z-scoring eigenvalues within respective GEPs. Z-scores for each gene were averaged across study to define the hnDAM signature (105 genes, average z-score > 5). (**C-D**) Expression of hnDAM genes across microglial subtypes shows enrichment in three DAM subtypes. Expression was computed using the *AverageExpression()* function in *Seurat*, and z-scored across subtypes (**C**), or a module score was assigned to each cell using the *AddModuleScore()* function (**D**). (**E**) Comparison of hnDAM to human^16,28,29^ and mouse^30,31^ AD-related microglial states highlight core drivers of disease-association. (**F**) GO enrichment of hnDAM signature is significantly enriched for lipid-related pathways (BH < 0.05) reduced using *rrvgo*^119^. See the supplementary information for a summary of all enrichment analyses (see **Table S34**). (**G**) Variance of hnDAM explained by expression of transcription factors within hnDAM (*MITF, PPARG, ARID5B, MAFB*) and putatively controlling microglia states^16,103,104^ (*CEBPA, SPI1*). Variance partitioning performed in *vegan*^102^ conditioning on the respective expression of other transcription factors (see **“hnDAM transcription factor analysis”**) with covariates for age and sex (*** FDR < 0.001, ** FDR < 0.01, * FDR < 0.05).

To summarize the GEPs associated with DAM-like subclusters, we selected the top 105 genes (average z-scored eigenvalue > 5) from Gerrits_2022_GEP5, Mathys_2024_GEP3, and Pineda_2024_GEP5 (**Fig. 3B**, **Table S33**), which included 97 protein-coding genes and 8 lncRNAs. The expression of this 105-gene hnDAM signature was largely specific to the DAM-like subclusters (**Fig. 3C**) and its average expression was increased in the 3 clusters of interest relative to all others (**Fig. 3D**). Compared to previously published human^16,28,29^ and mouse^30,31^ DAM signatures from AD and Aβ mouse models, the hnDAM signature shares key marker genes such as *GPNMB, SPP1, MYO1E* (**Fig. 3E**). By setting the z-score cutoff for GEP-derived eigenvalues to 5, we ensured that only those genes which most contribute to cross-NDD microglial activation were included in the final hnDAM signature. The hnDAM signature (**Table S7**) includes key hub genes that either modify the function of microglia in models of disease (e.g., *LGALS3*, *SPP1*)^32,33^ or carry genetic risk for one or more NDD (*ABCA1*, *ADARB1*, *DMXL2*, *DSCAM*, *FMN1*, *FNIP2*, *GPNMB*, *IQGAP2*, *LGALS3*, *MYO1D*, *NHSL1*, *PRKCE*, *RASGRP3*, *SAMD4A*, *SCARB2*, *SOCS6*, *SULT1C2*, *TANC2*, *UGCG*)^34–36^. Notably, *APOE* and *TREM2*, two AD risk genes commonly associated with DAM biology^37^, fell below our z-score threshold for inclusion in the final hnDAM signature, but were both positively associated with the consensus latent factor (*APOE*: z-score = 4.39, rank = 132, *TREM2*: z-score = 2.34, rank = 409, **Table S33**). This suggests that our hnDAM signature encapsulates a portion of previously published DAM signatures and markers, but adds genes in the context of multiple NDDs beyond amyloid-specific responses.

To further describe the pathways in which hnDAM genes are involved, we performed over-representation enrichment analysis followed by hierarchical organization summaries of GO:BP terms (**Fig. 3F**). We found that the hnDAM signature was significantly (FDR < 0.05) enriched for 42 pathways (**Table S34**). In agreement with previous work that has shown lipid regulation differences associated with the appearance of DAM-like cells^38,39^, the majority of significantly enriched terms were associated with “lipid localization,” “lipid catabolic process,” and “cholesterol transport.” This suggests that while the hnDAM signature is not specific to a single DAM-like subcluster, the biological themes previously reported in DAM-like cells are retained.

Previous work modeling transcription factor expression nominated *ARID5B*, *CEBPA*, *MITF*, *and PPARG* as putative regulators of DAM-like expression of a *GPNMB*-High microglia signature in AD^16^. In an effort to investigate how the hnDAM signature is regulated, we employed a similar approach to investigate the regulation of the hnDAM signature. The hnDAM signature contains four transcription factors: *ARID5B*, *MAFB*, *MITF*, and *PPARG*. The expression of these 4 transcription factors, alone and in combination, significantly explained the expression of the other 101 hnDAM genes in all discovery comparisons except Gerrits_2022_OC (**Fig. 3G**, **Table S35**). In comparison to the previous work^16^, three of these four hnDAM transcription factors (*ARID5B*, *MITF*, *PPARG*) were commonly nominated by both studies. However, hnDAM includes *MAFB* while excluding *CEBPA*, previously reported to control GPNMB^High^ DAM-like expression. Furthermore, *SPI1*, the gene encoding the transcription factor PU.1, did not explain much variability in the hnDAM signature except in the allocortex of AD and motor cortex of ALS/FTLD. Taken together, the four hnDAM transcription factors can influence the expression of the hnDAM signature in postmortem human cases of NDD.

Notably, the PD discovery cohort was excluded from this hnDAM signature definition since the study does not include the substantia nigra, where substantive neurodegeneration occurs. Most evidently, all four studies had one gene expression program specifically expressed in border-associated macrophages: Gerrits_2022_GEP9, Mathys_2024_GEP6, Pineda_2024_GEP6, NM_2024_GEP4 (**Fig. S10A**). Similar to our curation of hnDAMs, a consensus BAM factor was selected out of the top 149 genes (average z-scored eigenvalue > 5; **Fig. S10B**, **Table S36**). The expression of this 149-gene BAM signature was largely specific to the BAM subcluster (**Fig. S10C**) and its average expression was enriched in the BAM cluster, with some shared trends in monocytes and lymphocytes (**Fig. S10D**). This BAM factor shared 13 genes with the hnDAM factor (**Fig. S10E**), however the hnDAM genes appeared on a spectrum from negatively-associated to weakly-associated with BAM factor definition (**Fig. S10F**). These data suggest that while all four discovery datasets are sufficient for discovery of shared phenomena, the lack of a clear DAM gene expression program in the PD discovery cohort is not a limitation of the latent factorization pipeline.

### hnDAM stratifies donors along disease variables in discovery and replication datasets

Beyond the presence of clusters annotated as DAM-like, there is currently no standard test for whether DAM-like cells are present in a tissue. Since clustering and annotation are influenced by hyperparameter selection, large-scale replication across studies is challenging. To determine whether the hnDAM signature is replicable across other NDD cohorts, we implemented a two-step approach for validation. First, we performed a GSEA-based approach using the hnDAM signature to query lists of differentially expressed genes. Second, we used a PCA-based approach where the expression of hnDAM genes were used to score individual samples to test whether the hnDAM-score segregates donors by NDD.

Performing these tests in the discovery cohort, we demonstrate that the GSEA-based approach largely reflected microglial activation in AD and ALS/FTD spectrum cases (**Fig. 4A**, **Table S37**). In contrast, there was down-regulation of the hnDAM signature in the hippocampus of cognitive resilience and in the visual cortex of PD. Furthermore, the PCA-based approach was particularly robust for stratifying donors along the ALS/FTD spectrum. This was apparent in the motor cortex of FTLD patients (GSEA: NES = 2.44, FDR = 1.91E-14; PCA: β = 11.51, FDR = 4.28E-17; **Fig 4B**). Several regions such as the hippocampus of AD patients (GSEA: NES = 2.36, FDR = 3.71E-12; PCA: β = 4.10, FDR = 0.22; **Fig 4C**) and temporal cortex of FTD-GRN patients (GSEA: NES = 2.31, FDR = 5.50E-12; PCA: β = 5.61, FDR = 0.28; **Fig 4D**) displayed strong up-regulation of hnDAM genes. Finally, largely unaffected regions, such as the visual cortex of PD patients, displayed neither an up-regulation of hnDAM genes nor strong individual-level effects (GSEA: NES = -1.67, FDR = 0.0017; PCA: β = 0.65, FDR = 0.81; **Fig 4E**).

**Fig. 4.**
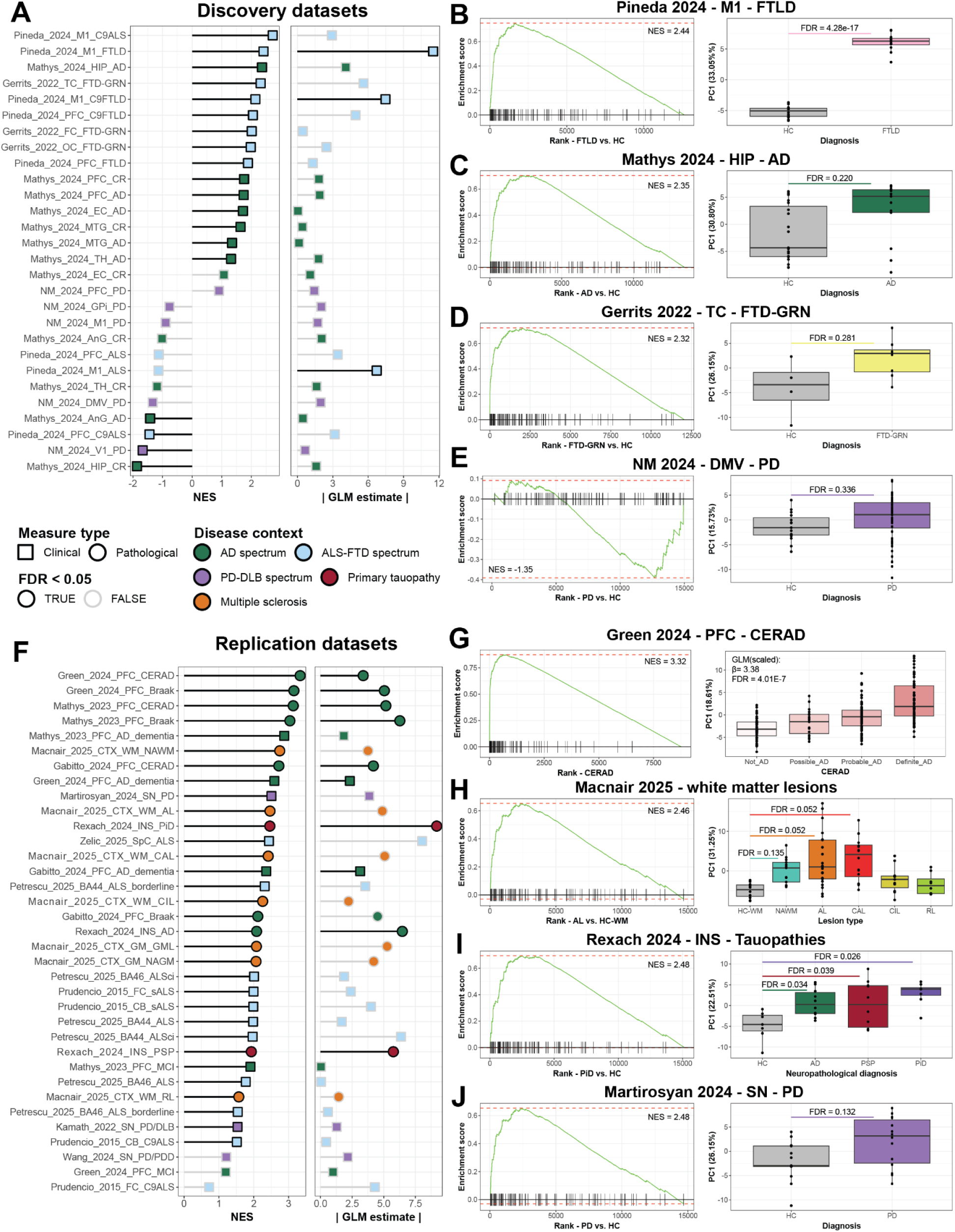
hnDAM stratifies donors by disease and pathology across discovery and replication cohorts. hnDAM signal was queried in independently collected sample series across NDDs (11 studies: **Table S38**) using complementary GSEA-based and PCA-based scoring (See **“Validation and replication of hnDAM signature”**). (**A**) Enrichment of hnDAM signature in the discovery datasets evaluated per comparison [disease x region] with GSEA-based (left) and PCA-based (right) methods. (**B-E**) Ranked expression of hnDAM genes and individual-level sample partitioning on hnDAM genes against NDD diagnosis. (**F**) Enrichment of hnDAM signature in the replication datasets evaluated per comparison [study x available disease variables] with GSEA-based (left) and PCA-based (right) methods.(**G-J**) Ranked expression of hnDAM genes and individual-level sample partitioning on hnDAM genes cognitive and neuropathological measures. P-value reported from *fgsea* and PCA-based GLM (*stats::glm()*) were Benjamini-Hochberg FDR-corrected within either discovery or replication cohorts. Statistics available in **Table S40**.

In order to demonstrate the replicability of the hnDAM signature and describe its presence across other NDDs and regions], we performed the GSEA- and PCA-based signature validation across 11 studies encompassing five age-related NDD spectra^15,40–49^ (AD, ALS/FTD, PD/DLB, primary tauopathy, and multiple sclerosis; see **Table S38** for replication cohort metadata). Making use of full-rank neuropathological information in the validation cohort, we were able to investigate whether the hnDAM signature correlates to a suite of neurodegeneration-associated clinical and pathological measures (for differential expression of replication datasets see **Table S39**). With this extra level of granularity, we observed a robustly up-regulated hnDAM signature via GSEA and individual-level stratification along both AD clinicopathological measures and types of primary tauopathy (**Fig. 4F**).

Notably, two of the highest NES scores for the hnDAM signature were found in differential expression models against CERAD classification (Green_2024: GSEA: NES = 3.33, FDR = 1.68E-32; PCA: β = 3.37, FDR = 4.01E-7; Mathys_2023: GSEA: NES = 3.13, FDR = 1.80E-24; PCA: β = 5.16, FDR = 9.97E-8; **Fig 4G, Table S40**). The strong linear correlation between hnDAM and an individual’s CERAD classification indicates that the appearance of cross-NDD microglial activation is closely linked to the progression of amyloidopathy. Notably, hnDAM was also significantly correlated with Braak stage (tauopathy progression) and AD dementia status (**Table S40**).

To bolster the claim that the appearance of hnDAMs are not unique to amyloidopathy, we included studies more broadly characterizing demyelinating disease and tauopathy. Evidence that the hnDAM signature is broadly generalizable across NDDs was especially striking in MS (Macnair_2025), where it was able to stratify samples from active lesions (AL) and chronically active lesions (CAL) compared to white matter from healthy controls (AL: GSEA: NES = 2.46, FDR = 2.73E-18; PCA: β = 4.89, FDR = 0.052; CAL: GSEA: NES = 2.43, FDR = 3.88E-17; PCA: β = 5.07, FDR = 0.052; **Fig 4H, Table S40**). Although these tests did not survive stringent FDR-correction across all 35 comparisons, the apparent PCA-based hnDAM increase in AL and CAL compared to unaffected controls and recovered MS lesions (i.e., CIL: chronically inactive lesion, RL: remyelinated lesion) suggests that the hnDAM signature can distinguish samples across clinically relevant measures in NDDs beyond ADRDs. Notably, by GSEA following differential expression compared to healthy tissue, the hnDAM signature was significantly enriched in all MS lesion types across white and grey matter (**Fig 4H, Table S40**).

Additionally, in a sample series of tauopathies (Rexach_2024), the hnDAM signature was strongly associated with tauopathy subtype and significantly stratified healthy controls from those diagnosed with AD, progressive supranuclear palsy (PSP), and Pick’s disease (PiD) (AD: GSEA: NES = 2.09, FDR = 1.43E-6; PCA: β = 6.47, FDR = 0.034; PSP: GSEA: NES = 1.94, FDR = 2.25E-5; PCA: β = 5.75, FDR = 0.039; PiD: GSEA: NES = 2.48, FDR = 1.8E-12; PCA: β = 9.19, FDR = 2.6E-3; **Fig 4I, Table S40**). This suggests that the hnDAM signature is increased in individuals with tau burden, regardless of clinical subtype or amyloid-β burden.

The discovery analysis of PD samples had no evidence for hnDAM up-regulation, and down-regulation in the visual cortex (**Fig. 4A**). To ask whether microglia respond with a hnDAM signature in the substantia nigra, the principal site of dopaminergic neurodegeneration in PD, we included three studies^42,46,49^ in our replication cohort which analyzed this region. In two of the three sample series, the hnDAM signature was significantly enriched via GSEA (Martirosyan_2024: NES = 2.48, FDR = 2.15E-11; Kamath_2022: NES = 1.56, FDR = 0.026; Wang_2024: NES = 1.23, FDR = 0.22; **Fig. 4J, Table S40**). However, expression of hnDAM genes was not sufficient to segregate patients from controls via PCA in any of the three studies (Martirosyan_2024: β = 3.86, FDR = 0.13; Kamath_2022: β = 1.28, FDR = 0.68; Wang_2024: β = 2.16, FDR = 0.38; **Figs. 4F, 4J, Table S40**). This suggests that while hnDAM expression may be present in PD, the magnitude, region, and timing of activation is more transient than other NDDs.

### Previously published chemical and biological perturbations induce partial hnDAM activation in iPSC-derived microglia

To develop an experimental model for the hnDAM signature, we used the SigCom LINCS database^50^ to identify chemical compounds which were predicted to mimic our hnDAM signature (**Fig. 5A**). This analysis identified 85 unique compounds (**Table S41**) predicted to induce hnDAM in four cell types (**Table S41**). From this list, we first chose to treat cells with the histone deacetylase inhibitor (HDACi) entinostat, along with a functional analog, vorinostat, and disease-associated molecular patterns from UV-exposed neurons (UV-DAMPs, see “**Production of Damage-Associated Molecular Patterns (DAMPs) from Neurons**”), all of which were previously shown to induce DAM-like states in mice^30^ and iPSC-derived microglia (iMicroglia)^51^.

**Fig. 5.**
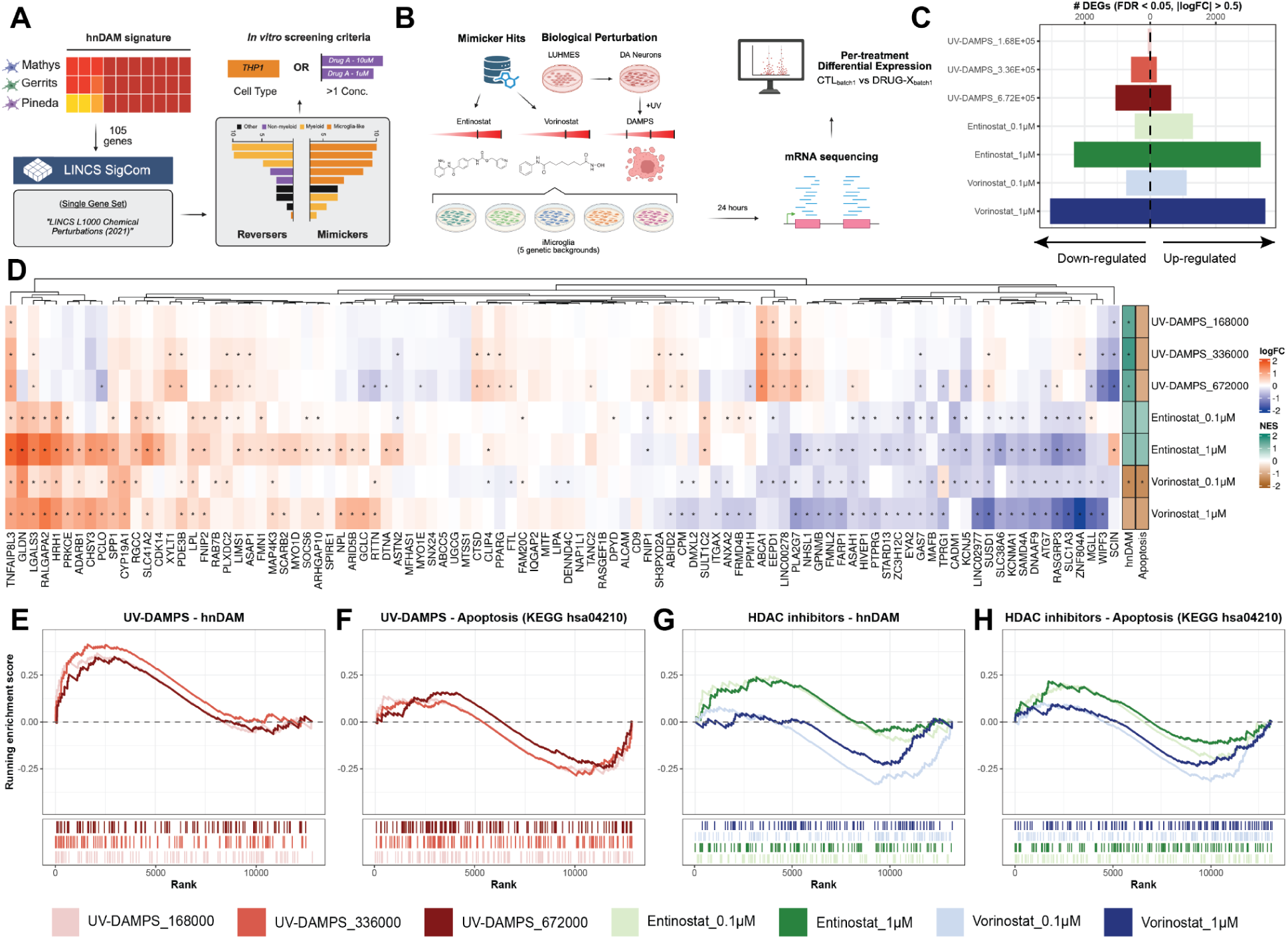
Chemical and biological perturbations induce aspects of hnDAM in iPSC-derived microglia. Transcriptional perturbations mimicking the hnDAM signature were predicted *in silico* using the Library of Integrated Network-based Cellular Signaling^50^ SigCom server (See **“Perturbation discovery for mimickers of hnDAM”**). (**A**) Schematic of perturbation discovery. (**B**) Experimental design for *in vitro* screening of perturbation. (**C**) Differential expressed gene sets of damage-associated molecular patterns (DAMPs) and HDAC inhibitor treatment of iPSC-derived microglia (n = 5 genetic background per condition). Analysis ran using *variancePartition::dream()* reveals dose-dependent effects in numbers of DEGs (|log2FC| > 0.5, FDR < 0.05). (**D**) Gene-level results for the 105-gene hnDAM signature reveal subsets recapitulated by either chemical or biological treatment. Enrichment for apoptosis (hsa04210) and hnDAM signature performed in *fgsea*. Significance stars for both DEGs and NES values indicate FDR < 0.05. (**E-H**) Ranked gene set plots from GSEA demonstrate normalized enrichment score from GSEA-calculations against hnDAM signature and apoptosis pathway. Statistics available in **Table S43**. Schematic (**A-B**) created with <u>BioRender.com</u>.

We identified differentially expressed genes across exposures using linear mixed modeling implemented through *dream*^52^. All three exposures exhibited dose-dependent effects, with UV-DAMPs having relatively milder transcriptional impacts compared to either HDACi at the doses tested (**Fig. 5B**). Of the 105 genes in the hnDAM signature, 98 were reliably detected in iMicroglia (**Fig. 5C**). From these, 87 were significantly differentially expressed (FDR < 0.05) in at least one condition, with 62 showing significant upregulation in at least one condition (**Fig. 5C**). Notably, hnDAM genes were not uniformly upregulated in response to either UV-DAMPs or HDAC inhibition. Between the two high-dose HDACi conditions (Entinostat_1μM and Vorinostat_1μM), 19 genes showed concordant upregulation, while 21 genes showed concordant downregulation (**Fig. 5C**). Between the two highest doses of UV-DAMPs treatment (UV-DAMPS_336000 and UV-DAMPS_672000), 17 genes were concordantly upregulated, while three were concordantly downregulated (**Fig. 5C**). Finally, across those four conditions, two genes (*TNFAIP8L3* and *LGALS3*) were concordantly upregulated, while one gene (*GAS7*) was concordantly downregulated (**Fig. 5C**). Using these differential expression results, we found that UV-DAMPs significantly induced the hnDAM signature without inducing apoptosis (**Fig. 5E-G**; **Table S43**). HDAC inhibition induced a partial, but not full, replication of the hnDAM signature (**Fig. 5G-H**; **Table S43**). The lack of apoptosis signatures in these treatments is expected as concentrations and timepoints were optimized for cell viability prior to transcriptional profiling (see **Methods**). The one exception was the 0.1-μM vorinostat treatment which we found significantly downregulated both the hnDAM signature (NES = -1.43, FDR = 0.041) and apoptosis (NES = -1.45, FDR = 0.041) relative to batch-respective controls. Taken together, this suggests that previously-published activators of DAM-like activation engage aspects of the hnDAM signature.

### PIKfyve and BTK inhibition induce robust hnDAM activation in iPSC-derived microglia

Among the compounds predicted to mimic the hnDAM signature were three additional kinase inhibitors: apilimod, a PIKfyve inhibitor, and spebrutinib and ibrutinib, both Burton’s tyrosine kinase (BTK) inhibitors (**Table S41**). We treated cells with these inhibitors each at multiple doses, along with another PIKfyve inhibitor, vacuolin. Additionally, as a control for non-hnDAM-related microglial activation, we also included exposures to interferon-gamma (IFNɣ), an inflammatory stimulus shown to up-regulate NDD-related genes such as *LRRK2* in microglia and peripheral immune cells^53,54^. Using an identical design to the previously published perturbations, we reliably detected 99 of the 105 hnDAM genes in a second batch of sequencing (**Figure 6A; Table S44**). Of these novel exposures, all but lower doses of ibrutinib exhibited robust, dose-dependent transcriptional impacts (**Fig. 6B**). Using these differential expression results, we found that both PIKfyve inhibitors and ibrutinib at all doses significantly induced the hnDAM signature without inducing apoptosis (**Fig. 6C; Table S45**). Notably, NES values for hnDAM induction via both PIKfyve inhibitors and ibrutinib were considerably higher than those for UV-DAMPs exposure (e.g., UV-DAMPS_336000: NES = 1.70, FDR = 0.00358; Apilimod_0.01μM: NES = 2.27, FDR = 2.09E-08; Vacuolin_0.1μM: NES = 2.44, FDR = 1.56E-09; Ibrutinib_0.1μM: NES = 2.26, 6.16E-09) (**Tables S43, S45**). In contrast, neither IFNɣ stimulation nor spebrutinib induced a statistically significant replication of the hnDAM signature. Taken together, our data suggests PIKfyve inhibition has the potential to robustly potentiate iPSC-derived microglia to a hnDAM-like state.

**Fig. 6.**
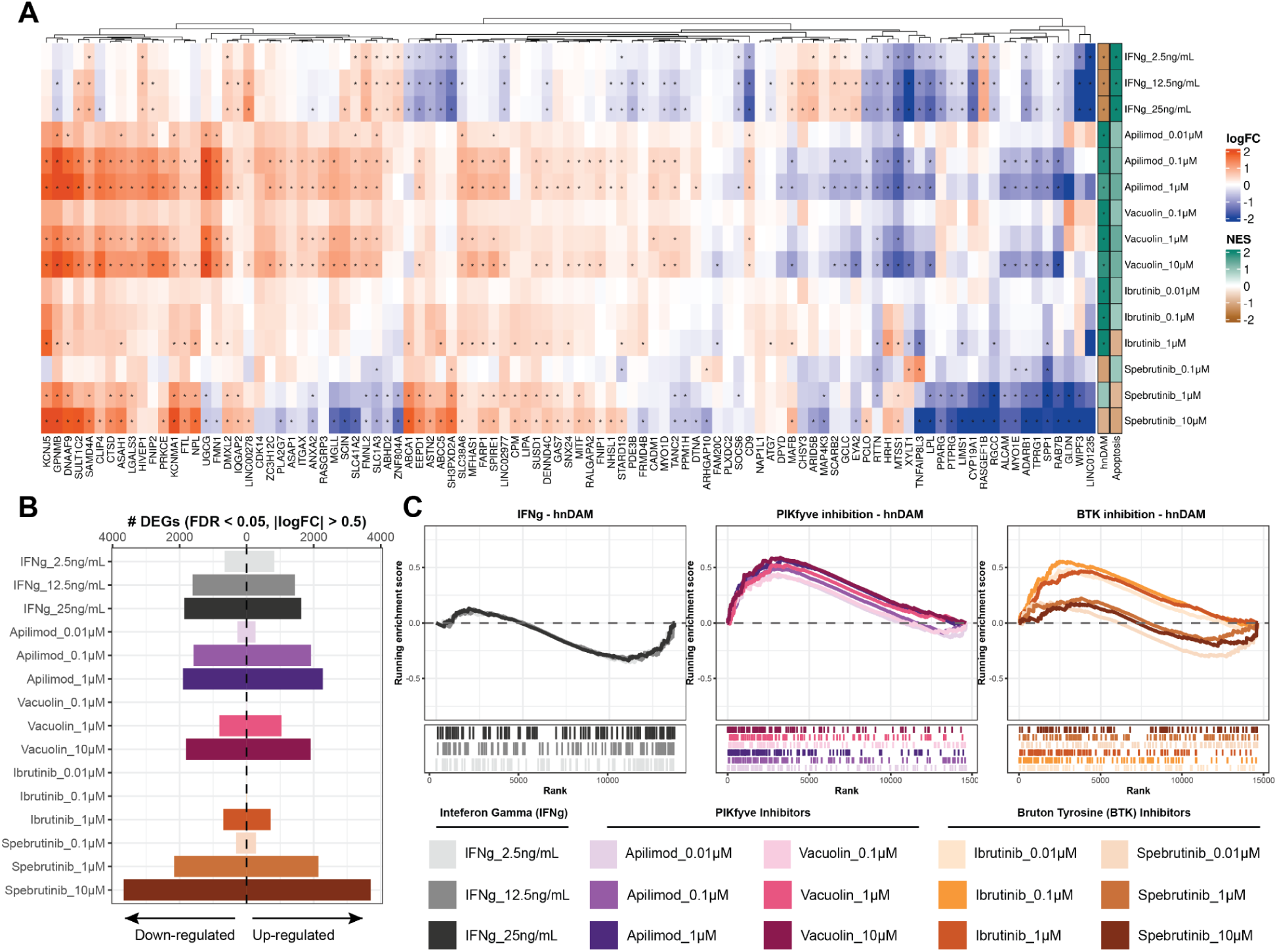
PIKfyve and BTK inhibition induce robust hnDAM activation in iPSC-derived microglia. Differentially expressed gene sets of IFNg stimulation, PIKfyve inhibition, and BTK inhibition of iPSC-derived microglia (n = 5 genetic background per condition). **(A)** Gene-level results for the 105-gene hnDAM signature reveal subsets recapitulated by novel mimickers. Significance stars for both DEGs and NES values indicate FDR < 0.05. (**B**) Robust transcriptional impact present across conditions (see **Table S44** for differential expression). Analysis ran using *variancePartition::dream()* reveals dose-dependent effects in numbers of DEGs (|log2FC| > 0.5, FDR < 0.05). (**C**) Ranked gene set plots from GSEA demonstrate normalized enrichment score from GSEA-calculations against hnDAM signature. Statistics available in **Table S45**.

Apilimod is an FDA-approved small molecule shown to have anti-proliferative effects in B-cell lines via IL-23 inhibition^55^, however any roles it might have in microglial function have not been demonstrated. In order to understand the transcriptional effects of PIKfyve inhibition via apilimod, we further characterized the differential expression results. We found a dose-dependent increase in the number of DEGs (**Fig. S11A**; 0.01μM = 535 DEGs, 0.1μM = 3,502 DEGs, 1μM = 4,170 DEGs) and the greatest overlap between 0.1μM and 1μM treatments (**Fig. S11B**; 3,244 shared DEGs, 92.63% of 0.1μM effects, 77.79% of 1μM effects). In an effort to discover what functions apilimod uniquely induces in microglia, we performed enrichment of the joint 0.1μM and 1μM apilimod effects but removed any genes overlapping with PIKfyve inhibitor analog vacuolin (**Fig. S11C**). We performed over-representation enrichment analysis followed by hierarchical organization summaries of GO:BP terms and found significant (B-H FDR < 0.05) enrichment for 37 pathways (**Table S46**). In agreement with previous work that has shown motility/migration effects of apilimod^56^, the highest of 13 summarized hierarchical terms included “myeloid leukocyte migration,” “extracellular structure organization,” and “regulation of taxis” (**Fig. S11D**, **Table S47**). This suggests that while the hnDAM signature is induced by apilimod, apilimod may also induce aspects of cell motility.

### hnDAM refines existing models of microglial activation

One challenge in the definition of DAM-like activation has been the lack of a testable readout. As a result, several studies have claimed aspects of DAM-like activation recapitulated *in vitro* based on single markers and/or limited functional assays^16,57^. In order to directly compare the effects of our transcriptional reprogramming of the hnDAM signature to existing studies of myeloid cell activation, we curated an *in vitro* replication cohort of antibody, cell debris, recombinant protein, and small molecule treatments across 12 studies (**Table S48**; Studies^17,51,57–65^). In comparison to the 12 of 22 treatments in the present study that were positively enriched for hnDAM via GSEA-based analysis, 15 of the 36 treatments in the *in vitro* replication cohort were similarly positively enriched for hnDAM (**Fig. S13, Table S49-50**). Notably these include: treatment with amyloid-β (Dolan_2023_Aβ, Smit_2022_Aβ), anti-TREM2 antibodies (Okuzono_2021_Hyb87), and HDAC inhibition (Haage_2025_Ent, Haage_2025_Vor). Although these studies varied in time, dose, myeloid cell model, and transcriptional assay (i.e. bulk vs. single-cell), hnDAM was significantly enriched in DEGs by hypergeometric test (FDR < 0.05, Fisher’s exact test) in 9 of the 12 treatments in this study and 6 of the 15 treatments in the *in vitro* replication cohort. 31 treatments (10 current study, 21 *in vitro* replication cohort) downregulated or failed to statistically modulate hnDAM. These studies vary in the number of biological replicates and overall signal recovered in differential expression. Where there were differentially expressed genes, hypergeometric analysis reveals which treatments were similar to and which were opposite of hnDAM activation in the human brain. Where there were not differentially expressed genes, GSEA-based analysis was still sensitive to describe the strong hnDAM responses to amyloid and novel pharmacological interventions. These complementary approaches offer directionality (sign NES), sensitivity (% hnDAM as DEG), and specificity (significance of hypergeometric test).

In addition to our integrative discovery cohort, several other studies have reported DAM-like signatures^16,30,31^. While direct comparison of gene sets can give a hint to the similarity of the activation types described (**Fig. 3E**), testing for the presence of effects mimicked in cell culture requires deeper characterization. We performed GSEA using the aforementioned gene sets against our *in vitro* replication cohort to investigate the efficiency of each set for evaluating activation profiles (**Fig. 13A-D, Table S50**). The strongest effects induced by debris (i.e. Dolan_2023_Aβ_iM, Dolan_2023_Apop_iM, UV-DAMPS) were captured by both human and mouse DAM-like signatures. However, the mouse DAM signatures resulted in divergent interpretations from the human signatures for TREM2 ASO treatment (Vandermeulen_2024_TREM2ASO-4W_iMXeno), BTK inhibition (Spebrutinib_1.00E-07) and Camptothecin (Tuddenham_2024_Camptothecin_HMC3). This suggests that microglia activation in the human brain is not only distinct from the responses of murine microglia, but also functionally mimicked by different compounds.

To determine whether sensitivity and specificity of microglia activation profiles are linked to causal microglia function in disease, we used a recent dataset of iPSC-derived microglia xenografted into the mouse brain treated with anti-amyloid antibodies^66^. To demonstrate the sensitivity of hnDAM (as defined) compared to other signatures and alternative hnDAM definitions with lower z-score thresholds, we performed GSEA on the results of Lecanemab-treated versus IgG control (**Fig. S14A, Tables S51-52**). The hnDAM signature and LDAM signatures (Haney_2024_AD_APOE44v33 and Haney_2024_iMG_APOE44_LDhi) were strongly induced by antibody treatment, while, mouse DAM-like signatures and the human GPNMB-high signature were not significantly enriched. However, the reported results relied on single-cell-level differential expression where there was a marked amount of gene sparsity (**Fig. S14B-C**; 43 of 105 hnDAM genes and 58 of 100 Marshe_2025_GPNMB.hi_human genes in results). Taken together, this suggests that the hnDAM signature is sensitive to identify microglia state changes that are not only responses to amyloid but associated with increased amyloid clearance.

## Discussion

In recent years, single-cell sequencing technologies have allowed deep molecular profiling of cell types in postmortem brains. Despite these technological advancements, the question of which disease-associated glial types are robustly present remains unanswered. By harmonizing astrocytes, microglia, and oligodendrocytes from four atlas-level multi-region NDD datasets, we demonstrate consensus substate annotations and test their disease association using cell type proportion and gene expression analyses (**Fig. 1**).We show that, once integrated, the atlases reveal directly comparable and disease-associated cell states that each single-disease study had not identified independently. We reveal prominent astrocytic regional heterogeneity, microglial contextual responses conserved across NDDs, and a lack of evidence for oligodendrocyte transcriptional diversity. This is consistent with findings in Alzheimer’s disease cohorts and mouse models of amyloidosis^18–21^, but is the first time the hypothesis has been systematically tested across NDDs and regions. We then define a consensus latent factor we termed “human neurodegenerative disease-associated microglia” (hnDAM) that captures cross-NDD microglia activation. We perform independent validation in a cohort containing 11 independent sample series across the NDD spectrum and *in vitro* using iPSC-derived microglia. Taken together, our data suggest that microglial activation to DAM-like states is not unique to Alzheimer’s disease and can be broadly generalized across NDDs.

First, we performed per-subcluster proportion analysis associating glial subtypes with NDD diagnoses across ten brain regions. Our analysis revealed that while astrocyte transcriptional identity was overwhelmingly dictated by brain region, microglia exhibited robust and transcriptionally similar disease-associated states that were conserved across NDD diagnoses. Our integrative framework uniquely enables discovery of AD-like DAM subclusters (e.g., Micro_DAM_GPNMB, Micro_DAM_SPP1) enriched in the brains of patients with TDP-43 proteinopathies. In contrast, oligodendrocytes displayed minimal heterogeneity related to either region or disease. This establishes that among glial cells, microglia are the primary indicators of conserved pan-neurodegenerative responses at the level of cellular subtypes (**Fig. 2**).

Glial activation has been viewed through simplified lenses such as pro- or anti-inflammatory microglia^5^ (M1 vs. M2) and neuro-toxic or -protective astrocytes^6^ (A1 vs. A2). Transcriptome-wide profiling enables a more nuanced, data-driven definition of contextual states based on gene expression. Our findings take a significant step towards this goal by demonstrating that despite drawing from multiple unrelated datasets, we can define a consensus disease association for microglia. This is further supported by microglial differential expression, revealing shared immune response pathways in microglia across diseases with deviations most apparent in unique contexts such as cognitive resilience to amyloid-driven dementia^19,44^.

While our data point to a dominant microglial response, this does not preclude the importance of disease-responsive gene expression profiles. While neither astrocyte nor oligodendrocyte substates were robustly associated with NDDs at the proportion level, large numbers of differentially expressed genes included a distinct downregulation of aerobic respiration pathways in astrocytes and oligodendrocytes of ALS patients. There also remains the possibility that these glial subtypes are responsive to pathology burden more than diagnosis status.

Having established that microglia mount a robust and conserved response at the level of proportions, we sought to move beyond this observation to create a quantitative definition of activation in NDDs. This led us to distill the core response into a unified signature that can be tracked, validated, and mechanistically interrogated. We calculated a core 105-gene latent factor, termed “hnDAM” (human neurodegenerative disease-associated microglia; **Fig. 3**). The signature was commonly attributable to 3 DAM-like subclusters in our discovery cohort and found to be replicated across 11 independent human postmortem brain datasets, encompassing a wide spectrum of NDD pathologies including Alzheimer’s disease, synucleinopathies, TDP-43 proteinopathies, primary tauopathies, and demyelinating disease (**Fig. 4**). The hnDAM signature offers a unifying quantitative framework to track a specific, conserved microglial activation state across the landscape of human neurodegeneration. Interestingly, the hnDAM signature includes transcription factors previously hypothesized to control DAM-like induction^33,38^ and 19 GWAS candidates from AD, PD, and ALS studies^34–36^.

Understanding of microglial activation has been shaped by work in mouse models, which first described “Disease-Associated Microglia”^31^ and “Neurodegenerative Microglia”^30^ in the context of amyloidosis. These studies establish a powerful paradigm: that glial cells adopt context-specific but plastic transcriptional states in response to neurodegenerative insults. While some studies from postmortem AD tissue identify disease-associated populations similar to the murine DAM signature^16,28,29^, the signature’s existence and composition are hotly debated. Gene lists used to determine DAM presence are often derived from subsets of markers and prioritize individual genes over their combined action. Having established the hnDAM signature as a robust and replicable marker of the microglia contextual state shared across NDDs, we sought to develop a tractable *in vitro* model for further investigation.

The goal of modeling reactive microglial states *in vitro* has been a major focus for the field, with several groups making critical advances using various approaches. These include perturbation screens, targeted genetic manipulations, and combinatorial screens of differentiation factors^38,51,57^. However, a persistent challenge has been the lack of a readout for the targeted contextual state, with most studies benchmarking against the murine DAM signature or *ad hoc* markers that do not generalize. Our work builds directly upon these foundational efforts but provides a key refinement: we use our systematically defined and validated hnDAM signature from human brains as a statistically-testable endpoint.

Transcriptional reprogramming prediction is a burgeoning new strategy employed in several high impact publications to repurpose off-target effects of approved drugs in potentially beneficial contexts^67–70^. After predicting compounds that would mimic the hnDAM signature *in silico*, we treated iMicroglia with several sub-toxic doses for 24 hours *in vitro*. We recognize that treatments such as IFNɣ and exposure to neuronal debris have been reported to induce microglial activation and included them as positive controls. We found that only a subset of the hnDAM signature genes were recapitulated when treated with chemical or biological perturbations previously reported to induce activation in iPSC-derived microglia (**Fig. 5**). Notably, exposure to neuronal debris (UV-DAMPs) served as a potent stimulus of lipid-related features without triggering apoptosis. In contrast, histone deacetylase inhibitors (HDACi) modulated expression of hnDAM genes, upregulating 19 genes while downregulating 21. Both previously demonstrated perturbations failed to robustly induce holistic DAM-like signatures, however in our hands, neuronal debris and HDACi induced complementary aspects of hnDAM.

We additionally demonstrate that two PIKfyve inhibitors, apilimod and vacuolin, both induce hnDAM expression more robustly than either UV-DAMPs or HDACi (**Fig. 6**). Apilimod is in phase 2 clinical trials for both SARS-CoV2 and C9orf72-linked ALS^71^, where it increased soluble plasma GPNMB by >2.5-fold while lowering CSF poly(GP) (an ALS biomarker) by 73%^72^. Conversely, only one of the two BTK inhibitors identified by LINCS, ibrutinib, significantly induced hnDAM, while spebrutinib did not. Additionally, alternative microglial activation via IFNɣ stimulation did not induce hnDAM-related gene expression. These data demonstrate that neither type-II interferon activation nor BTK inhibition were sufficient to engage the transcriptional signature we found across discovery and replication NDD cohorts.

Importantly, our *in silico* analysis nominated other mimickers of the 105-gene signature based on transcriptional reprogramming prediction from LINCS^50^, opening the door to high-throughput screens to identify novel therapeutics that can either prevent or induce hnDAM.

Unified discovery across NDDs pinpoint microglia as principal effectors of glial activation shared across the degenerating human brain. From a clinical perspective, our data suggests the hnDAM signature could be broadly applied across multiple disorders. Importantly, this would offer a generalizable strategy for treating a growing population of patients presenting with co-pathologies. By applying statistical testing to the nature of reactive glia, disease associations as we define them extend beyond Alzheimer’s disease and models of amyloidosis. While disease-specific patterns of activation may still exist both in endpoint studies and during disease progression, the unifying hnDAM signature outperforms existing signatures in strongly recognizing disease-associated activation when it is present, reliably evaluating the efficiency of cell culture models, while still capturing activation that is casually involved in the removal of amyloid plaques.

We acknowledge several limitations to our study. Our integrative approach, while allowing for the discovery of generalizable glial substates, relies on combining previously published datasets which introduces dataset limitations intrinsic to the specific hypotheses originally tested (e.g., region, age, sex). Similarly, since neuropathology is reported in a manner specific to each primary study, full rank neuropathological scoring of all samples from all regions was not possible. Future studies with harmonized neuropathology across NDDs will be necessary to resolve other transient cell states and determine the causality of hnDAMs in the complex landscape of neurodegeneration. Additionally, the use of iPSC-derived cell culture models do not fully recapitulate microglia states in the human brain, however our analysis demonstrates their flexibility when treated with biologically-informed exposures. Therefore, our work provides a critical foundation and a tangible molecular signature for the development of pan-neurodegenerative therapeutics targeting microglia.

## Methodology

### snRNA-seq data preprocessing and quality control

Fastq files were downloaded from their respective sources (see Data Availability). Reads were mapped and cell-by-gene count matrices were generated using *CellRanger* (10x Genomics, v8.0.1), with the 2024-A reference transcriptome. To correct for ambient RNA contamination and identify empty droplets, we used *CellBender*^73^ with the default settings (learning rate = 1e-4, epochs = 150). To identify doublets, we ran the packages *scds*^74^, *scDblFinder*^75^, and *DoubletDetection*^76^, all wrapped in the *Demuxafy* singularity container^77^. We removed nuclei which at least 2 of the 3 doublet detection algorithms called as doublets. For genotype-based demultiplexing of nuclei from different donors run in the same 10x lane (applicable only to the NM_2024 dataset), we utilized the packages *Demuxalot*^78^ and *Cellsnp-lite/Vireo*^79,80^, again wrapped by *Demuxafy*. We removed nuclei which either demultiplexing software called as a heterogenic doublet, along with any nuclei for which the 2 softwares did not assign the same donor. Finally, we performed cell-level quality control (QC) for each dataset independently, removing nuclei which had UMI counts in the top or bottom 2.5% or UMI features in the bottom 2.5% for their respective dataset, as well as nuclei with greater than 2% mitochondrial or ribosomal counts. Note that doublet detection and demultiplexing algorithms, as well as cell-level QC filtering, were performed on the uncorrected *CellRanger* counts matrices, not *CellBender* ambient-corrected counts. Cells that passed all QC filters were retained and we used their respective *CellBender*-corrected counts for all downstream analysis (**Table S1**).

### Initial processing of individual snRNA-seq datasets

Samples from the various datasets were first merged within their respective dataset by brain region, and the standard *Seurat*^22^ pipeline was performed (*NormalizeData()*, *FindVariableFeatures()*, *ScaleData()*, *RunPCA()*) with batch integration using *Harmony*^81^. *FindNeighbors()* was run using an appropriate number of dimensions based on the *Harmony*-corrected principal components, and broad clusters were computed using *FindClusters()* at a resolution of 0.1. Following this initial clustering, microglia, astrocytes, and oligodendrocytes were identified based on expression of canonical marker genes (*CSF1R*, *AQP4/GFAP*, and *MOG*, respectively), and subset out of the main objects. Next, for each dataset independently, microglia, astrocytes, and oligodendrocytes (each cell type separately) from each brain region were merged and processed in the same manner as previously. We manually removed clusters which co-expressed canonical marker genes of another cell type, or, in the case of microglia subsets, expressed markers of T-cells or other peripheral immune cells (e.g., *CD3E*). Finally, any [donor x region] which did not have at least 50 cells of a respective cell type was removed.

### Atlas-scale integration and clustering of snRNA-seq datasets

Due to large differences in both sequencing depth and number of cells provided from each dataset and based on extensive testing of different integration options, we opted to use *Seurat’s* sketch-based integration workflow. Briefly, this process assigns each cell a “leverage score”, corresponding to the relative amount of variance that cell explains within the context of a unit of data (in our case, [dataset x region]), and selects a set number of cells from each unit of data, prioritizing cells with higher leverage scores (i.e., those cells that contribute the largest amount of variance)^22^. We chose this option to mitigate the eventuality we observed in testing whereby, for example, the Gerrits_2022 study, which contributed considerably more cells per donor than the other datasets due to their flow-sorting strategy, while having lower numbers of UMIs and unique genes per nucleus, would dominate the clustering, leading to subclusters and marker genes which simply recapitulated those found in that individual study. For this approach, we “sketched” a number of cells per [dataset x region] which was slightly lower than the lowest number of cells provided by a single [dataset x region] (microglia: 6,000; subcortical astrocytes: 20,000; cortical astrocytes: 13,000; oligodendrocytes: 30,000). Next, we performed the same *Seurat* workflow as described previously on the sketched cells, including *FindVariableFeatures()*, *ScaleData()*, and *RunPCA()*, *Harmony* batch correction across dataset and donor, followed by *FindNeighbors()* and *FindClusters()*. For determining optimal clustering resolution, we clustered at a range of resolutions, ranging from 0.05 to 0.5. We chose the resolution to proceed with based on prior biological knowledge (including both evaluating subcluster marker genes at different resolutions and comparing to published signatures; see **“Characterization of cell subtypes”** below) and cluster stability, evaluated using the *clustree* package^82^. The resolutions we proceeded with were as follows: microglia: 0.25; all astrocytes: 0.15; subcortical astrocytes: 0.2; cortical astrocytes: 0.15; oligodendrocytes: 0.15. Once we decided on a clustering resolution, we re-projected the remaining cells into the *Harmony*-corrected latent space of the sketched cells using *ProjectIntegration()*, transferred cell subtype labels using *ProjectData()*, and re-computed the UMAP with *RunUMAP()*, all using the same number of dimensions determined from the *Harmony* principal components. During clustering, some additional clusters of low-quality nuclei (relatively low UMI counts or high mitochondrial reads) or suspected doublets were identified and removed.

### Characterization of cell subtypes

Marker genes for cell subtypes were computed using the *FindAllMarkers()* function in *Seurat*, with the only filter being the proportion of cells in a cluster expressing a gene >= 0.2. For comparison to published gene signatures or marker lists, we used gene set enrichment analysis^83^ implemented through the *fgsea* package^84^ in R, setting eps = 0 and nPermSimple = 10000. Genes used for comparison from published sources can be found in **Tables S3-5**^14,16,28–31,41,51,85–89^.

### Cell subtype proportion analysis

To describe the effect of region on cell subtypes, cell counts were calculated aggregated by region. The effect of region class (cortical and subcortical) was tested via Welch’s t-test and reported as nominal p-values.

To investigate the effect of disease status on glial subtypes, cell counts were aggregated by [donor x region], and proportions were calculated by dividing the number of cells of a given subtype by the total number of cells from that [donor x region]. Given that proportion values are *de facto* compressed from 0-1, we applied a logit transformation; prior to transformation, we added and subtracted an offset of 1e-6 to proportion values of 0 and 1, respectively. To test differences in cell type proportions by disease group, we applied a GLM with the formula proportion ∼ disease group + sex + scaled age. Note that we chose to perform these comparisons in a “1 vs. 1” manner, between healthy control donors and donors from a single disease category from a given [dataset x region]. Additionally, since public ROSMAP age reporting is cut off at 89, and binning was not possible due to limited sample sizes, all ages 90+ were set to 90 for the Mathys_2024 study. P-values from GLM models were corrected using the Benjamini-Hochberg method.

We then applied the R package *mashr*^90^ for meta-analysis of proportions data, which examines patterns of similarity across conditions to estimate and refine effect sizes, their directions, and p-values. In our case, we input the GLM estimates and standard errors from proportions comparisons (disease vs. HC in a given [dataset x brain region]) to *mashr*, computed canonical covariance matrices, selected a subset of strong signals at a local false sign rate (*LFSR*) threshold of 0.05, and when possible, identified data-driven patterns of covariance across the proportions data. For *mashr* runs where no comparisons met the *LFSR* threshold for the strong subset (i.e., cortical astrocytes and oligodendrocytes), canonical covariance matrices and their associated β_mashr_ and *LFSR* values were used for visualization and interpretation. β_mashr_ and *LFSR* values reported for microglia tests are refined values derived from data-driven covariance analysis.

### Differential gene expression analysis

We performed differential expression (DE) analysis using *DESeq2*^91^ for a pseudobulk approach, and *nebula*^92^, which uses a negative binomial mixed model at the single-cell level. For both approaches, we prepared the data in the same manner. Each DE comparison was performed within a single [dataset x region], using healthy control or diseased donors from that dataset, comparing healthy controls to a single disease category at a time (if a given study had multiple). First, for a given [dataset x region x disease] comparison, we filtered raw single-cell counts matrices for genes which were expressed in at least 0.5% of all cells, as recommended by *nebula*. Counts were then aggregated by donor.

Since biological and technical metadata for the different studies was not full-rank (e.g., ROSMAP metadata for age cuts off at 90, missing postmortem interval information across studies, etc.), and we did not use different covariates across different datasets, we opted to use an approach for removal of unwanted variance using the *RUVseq* package in R^93^, similar the that taken in Mathys_2024^19^. For this, we first TMM-normalized^94^ the donor-pseudobulked counts and ran a GLM simply with the design formula GEx ∼ disease group. We then extracted the residuals (i.e., all variance not associated with disease group) from the GLM and ran *RUVseq* on them (with the *RUVr()* flavor, for identifying components of unwanted variance from residuals), with the *k* (number of components of unwanted variance to find) set to the number of donors in that comparison / 2. Next, we determined the optimal number of components of unwanted variance to use as covariates in the DE analysis in an unsupervised manner. We used the *variancePartition* package^95^ to iteratively fit models, adding an additional component of unwanted variance to the design each time (i.e., ∼ disease group + component_1 + component_2 … + component_n). After each iteration, we extracted the value for the percent of total variance explained by the residual. After all components had been added, we used the *ede()* function in the R package *inflection*^96^ to determine the maximal inflection point of the elbow plot generated from the percent variance explained by the residual from each iteration of *variancePartition*. This served as the number of components of unwanted variation to add to the final design matrices for DE analysis.

For DE analysis, we input either the pseudobulked (for *DESeq2*) or single-cell (for *nebula*) counts matrices, along with donor information and values for components of unwanted variance, with the design formula GEx ∼ disease group + component_1 … + component_n. We ran *DESeq2* using the standard settings, and for *nebula*, we used cells’ number of UMIs (nCount_RNA) as the offset, donor as the random effect variable (both as recommended), and added percent mitochondrial UMIs per cell to the design formula. For downstream analysis, we only considered genes for which *DESeq2* did not produce *NA*s for p-value/padj, convergence in the *nebula* analysis was equal to 1, and the standard error in the *nebula* analysis was less than 10. To confirm the robustness of our DE results across analysis methods (pseudobulk vs. single-cell), we also regressed the beta (i.e., log_2_FC) values from both methods for each gene against each other (**Fig. S5D-E**), and found strong correlation between the two methods across datasets and comparisons. Given this, we proceeded with downstream analysis using values from the *nebula* analysis.

### Meta-analysis of differential expression results with *mashr*

Since log_2_FC values (disease vs. control) across comparisons were not directly comparable (due to different sample sizes, numbers of cells, etc.), we z-scored log_2_FC values within each comparison and used z-scores and non-adjusted p-values (both from *nebula* results) as inputs for *mashr*. Similar to the proportions meta-analysis, we began by generating canonical covariance matrices (*cov_canonical()* function), and set up the main data object with this correlation structure (*mash_set_data()* function). We then selected a strong subset of signals using a condition-by-condition analysis (*mash_1by1()* function); to limit dataset effects in the meta-analysis, we used a stringent *LFSR* threshold of 1e-4 for selecting this strong subset of genes. We then performed data-driven covariance analysis by first running PCA on the strong subset (*cov_pca()* function, using all PCs that explained greater than 5% overall variance), then using those matrices to run extreme deconvolution (*cov_ed()* function)^90^. Finally, we once again fit mash with both the canonical and data-driven covariance matrices, and extracted the fitted g mixture from this model.

Following *mashr* analysis, we examined the estimated mixture proportions (*get_estimated_pi()* function) to find which data-driven covariance matrices were the most common pattern in the data. We then extracted a list of genes whose patterns of variance were best fit to the “ED_tPCA” data-driven covariance matrix (i.e., pattern). Using the *mashr*-refined beta values for these genes, we *k*-means clustered them, determining an optimal *k* by examining the total within-cluster sum of squares at values of *k* ranging from 1 to 10. These *k*-means clusters are presented in the heatmaps in **Figs. S6-8** and in **Tables S19, S23, and S27**. For given *k*-means clusters of interest, we performed functional enrichment analysis on the genes within using the *ClusterProfiler* package in R^97^, testing against the Gene Ontology: Biological Process term set^98,99^. We present the top five most significantly enriched pathways for clusters of interest in **Figs. S6-8**.

### Microglia signature discovery via latent factorization

Latent factorization for the purpose of module discovery was performed using consensus non-negative matrix factorization (*cNMF*)^100^. Given differences in sequencing depth between datasets and our desire to independently discover an equivalent DAM gene expression module in multiple datasets, we chose to run *cNMF* on the discovery datasets separately. We first preprocessed the data for *cNMF* by randomly downsampling each dataset to 25,000 nuclei, importing raw single-cell counts from each dataset into Python, and running the *preprocess_for_cnmf()* function, which library-size normalizes raw counts, finds highly variable features, and performs batch correction. We then prepared and factorized the data, running *cNMF* at values of *k* ranging from 5 to 35, with 100 iterations per *k*. The optimal *k* value was manually chosen for each dataset by balancing high cluster stability and low error rate (Gerrits_2022 *k* = 16; Pineda_2024 *k* = 12; Mathys_2024 *k* = 12, NM_2024 *k* = 17, **Fig. S9**). After final *k*-clustering was complete, we inspected the GEP usage per cell averaged across subclusters. We observed that in three datasets, there was a GEP (Gerrits: GEP5; Pineda: GEP5; Mathys: GEP3) which was enriched in all 3 of Micro_DAM_SPP1, Micro_DAM_Int2, and Micro_DAM_GPNMB clusters relative to other subclusters; this GEP in each dataset was considered the “hnDAM” module. We also observed that across all four datasets, there was a GEP (Gerrits: GEP9; Pineda: GEP6; Mathys: GEP6, NM: GEP4) which was enriched in BAM clusters relative to other subclusters; this GEP in each dataset was considered the “BAM” module. For defining the consensus signatures across datasets, we first filtered for genes which were detected in all 3 datasets, and z-scored the contribution (i.e., eigenvalue) of each gene to each of the respective dataset modules of interest. We then averaged the z-scores of each gene’s contribution across datasets, and defined consensus modules as having an average z-score contribution of at least 5.

For evaluating pathways enriched in our hnDAM signature, we used the R package *ClusterProfiler* to obtain GO:BP terms, then used the package *rrvgo* to group and simplify the terms with a Benjamini-Hochberg-corrected FDR < 0.05.

### Validation and replication of hnDAM signature

We tested the predictive power of our hnDAM signature in our 4 discovery datasets (Mathys 2024, NM 2024, Pineda 2024, Gerrits 2024) and 11 replication datasets (**Table S38**)^15,40–49^ using two complementary methods. For replication datasets, we only used available processed data from the studies, which was downloaded from the respective sources (GEO, Synapse, etc.). Note that where fully processed objects with annotations were available, we used the authors’ original annotations of microglia. However, some studies did not have this available, in which case, we integrated samples, performed broad clustering, and identified microglia based on canonical marker gene expression (i.e., *CSF1R*). Notably two of the studies in the replication also included donors from the ROSMAP cohort. In order to avoid pseudo-replication at the level of donor: any individuals from Mathys_2024 (discovery dataset, n = 48) were removed from other ROSMAP studies (Green_2024 and Mathys_2023), any overlapping donors between the remaining studies were assigned to the older dataset (Mathys_2023, n = 315), and the newest dataset was subset down to only the unique donors (Green_2024, n = 217). All other replication datasets were used as provided by their respective studies.

First, we asked whether the hnDAM signature could segregate donors based on biological disease variables (i.e., disease status, neuropathology, etc.). To accomplish this, we developed a summary-scoring method for the hnDAM signature (referred to as “PCA-based analysis”). First, counts were aggregated across [donor x region] and TMM-normalized using the *edgeR* package. Next, pseudobulk counts matrices were filtered to only include genes in the hnDAM signature. Note that, since we used only processed data from replication datasets, some hnDAM genes may have been missing. We also removed any genes which had 0 counts in all samples (and thus had 0 variance). We then performed PCA analysis on the normalized hnDAM counts, and ran a GLM with the formula PC1 ∼ disease_variable + covariates, where PC1 was the value of the first principal component for each donor, disease_variable was the biological disease variable of interest, and covariates included whatever metadata was available for the samples (e.g., sex, age, postmortem interval, brain bank, etc.). For the discovery cohort, we used sex and age as covariates. P-values obtained from the GLMs were FDR-corrected across the discovery or replication cohorts. Finally, note that for the replication cohort, since they did not undergo an identical preprocessing pipeline to the discovery cohort, we performed the Rosner test for outliers on the values of PC1, removed any samples identified as outliers, and re-computed the PCA; this process was repeated until no outliers remained.

Since this PCA-based analysis is inherently directionless (i.e., the value of PC1 between disease and control does not indicate enrichment or depletion of the hnDAM signature, only separation of donors by expression of those genes), we performed a complementary gene-set enrichment analysis (GSEA) in the discovery and replication datasets (referred to as “GSEA-based analysis”). For the discovery datasets, we used the differential expression results from *nebula* for each comparison. For the replication datasets, we performed differential expression analysis using a pseudobulk approach with *DESeq2*. Briefly, microglial single-cell counts were aggregated by donor, and we ran *DESeq* with the default settings, using terms in the design formula for available covariate metadata (the same covariates used in the GLM for PCA-based analysis). Next, for both discovery and replication datasets, we ranked genes in the differential expression analysis by the signed -log_10_(p-value) (i.e., sign(log_2_FC) * -log_10_(p-value)), and performed GSEA as implemented through the *fgsea* package in R against our hnDAM signature. We FDR-corrected the p-values from discovery and replication datasets, and report normalized enrichment scores (NES) and FDRs.

### hnDAM transcription factor analysis

To determine the contribution of different transcription factors (TFs) to driving the hnDAM gene expression signature, we took a variance partitioning approach. We first identified which genes in the hnDAM signature were themselves TFs by comparing the 105-gene list to the HOCOMOCOv11 TF database^101^, which identified *MITF*, *PPARG*, *ARID5B*, and *MAFB*.

To determine the contribution of each TF to expression of the overall hnDAM signature, we first summarized hnDAM expression by [donor x region] using the same approach as described above, but first removing the 4 TFs from the counts matrix prior to PCA (note that this and subsequent variance partitioning analysis was performed by [dataset x region], agnostic of disease group). To remove the influence of covariates, we performed a GLM with the formula PC1 ∼ age + sex and extracted the residuals (i.e., other variance not associated with those variables). We then used the *varpart* function from the R package *vegan*^102^ to assess the variance explained by the expression of each of the 4 TFs on the rest of the hnDAM signature. Briefly, we input the residuals from the GLM as the test variable, and scaled expression of the four TFs as explanatory variables. From this model, we extracted several outputs: the variance explained by each of the 4 TFs when conditioned on the expression of the other 3 TFs (“variance uniquely explained”), and the total variance explained by the expression of all 4 TFs in combination. To obtain p-values for these contributions, we used the *anova.cca* function from the *vegan* package. For testing the variance uniquely explained by each TF, we conditioned on the expression of the other 3 TFs and covariates (e.g., formula = PC1 ∼ *MITF* + condition(*PPARG* + *ARID5B* + *MAFB* + age + sex)), and for the combinatorial variance explained, we only conditioned on covariates (e.g., formula = residualized PC1 ∼ *MITF* + *PPARG* + *ARID5B* + *MAFB* + condition(age + sex)).

To evaluate the specificity of the variance of hnDAM signature explained by the four TFs, we tested the variance explained by two other TFs with known involvement in microglial disease biology, PU.1 (*SPI1*) and *CEBPA*^16,103,104^. Here, we tested the unique variance explained by the expression of these two TFs conditioned on the expression of hnDAM TFs as above (e.g., formula = PC1 ∼ *SPI1* + condition(*MITF* + *PPARG* + *ARID5B* + *MAFB* + age + sex)).

Note that due to the nature of variance partitioning with *vegan*, values for proportion of variance explained will occasionally be negative; this occurs when the explanatory variable being tested explains less variance than a random variable would, and these instances are to be interpreted as 0 variance explained. For any such instances in our data, we set the variance explained to 0 and removed the p-value prior to FDR correction. All p-values were FDR-corrected using the Benjamini-Hochberg method.

### Perturbation discovery for mimickers of hnDAM

To determine chemical perturbations that are predicted to mimic the hnDAM signature, we employed the Signature Commons (SigCom) search from the Library of Integrated Network-based Cellular Signatures (LINCS) program^50^. In brief, the search function uses previously computed gene rankings of L1000 signatures^105^ and runs a Mann-Whitney *U* test on up- and down-regulated gene sets to calculate z-score. We input the 105-gene hnDAM signature as a single gene set in the server and assessed which treatments were scored as the best mimickers, using the “LINCS L1000 Chemical Perturbations (2021)” database. Perturbations were further prioritized based on cell line assayed and “positive mimic” hits from drugs at multiple concentrations.

### Differentiation of iPSC-derived Microglia

Human induced pluripotent stem cells (iPSCs) were obtained from the Michael J. Fox Foundation’s Parkinson’s Progression Markers Initiative^106^ (PPMI). Five clonal lines from healthy donors with no known neurodegeneration risk (confirmed not to have pathogenic variants within SNCA, LRRK2, or GBA1) were selected as biological replicates for the entire study (PPMI-3453, PPMI-3658, PPMI-3480, PPMI-3668, PPMI-3460). iPSC-derived microglia (iMicroglia) were differentiated via embryoid-body mediated protocol, with the following specified modifications. In brief, cryo-preserved iPSCs were plated in StemFlex^TM^ Basal media (Gibco #A33493-01) in 6-well plates coated with 167 μg/mL Matrigel® Matrix (Corning #354277 diluted in in DMEM/F12 (Gibco #11320-033)) and fed every other day until approximately 60-70% confluent. To start differentiation, the iPSCs were formed into embryoid bodies by seeding 10,000 cells in each well of a 96-well v-bottom plate in 100uL of E8 media (Gibco #A1517001) with 100 ng/mL VEGF-121 (ThermoFisher #100-20A) and BMP-4 (ThermoFisher #120-05ET), 20 ng/mL SCF (ThermoFisher #300-07), and ROCK inhibitor (STEMCELL Technologies # Y-27632). Embryoid-body media changes were performed daily for 3 days, removing the ROCK inhibitor after the first 24 hours. To begin the production of primitive macrophage precursors (PMPs), approximately 400 embryoid bodies were transferred into a 175 cm^2^ flask coated with 2 μg/mL of fibronectin (Invitrogen #33016015) on day 4. Microglia progenitor media (X-Vivo medium (LONZA #00011355250) with 2 mM GlutaMax (Gibco #35050061), 55mM beta-Mercaptoethanol (Gibco #21985023), 100ng/mL M-CSF (ThermoFisher #300-25), and 25 ng/mL IL-3 (ThermoFisher #200-03)) was changed every other day until release of PMPs by embryoid bodies into the media was apparent (10-21 days depending on differentiation). PMP identity was determined by attachment. First, cells were centrifuged at 300g for 3 minutes and plating 600,000 cells into a single well of a 6-well plate. If PMPs did not attach to uncoated plastic wells within 2 hours, cells were determined to not be mature enough to be collected for the final step of maturation. Embryoid bodies producing PMPs were maintained for up to 4 months and collections performed biweekly.

To complete iMicroglia maturation, once production of microglia progenitors began sometime from day 10-21 in the microglia progenitor media and attachment test was successful, progenitors were transferred every twice per week into 6-well plates coated with 50 μg/mL Poly-L-orinthine (Sigma-Aldrich #P4957) and 1 μg/mL fibronectin, where microglia maturation media, Advanced RPMI (Gibco #12633012) with 2 mM GlutaMax (Gibco #35050061), 100 ng/mL IL-34 (ThermoFisher #200-34), and 10 ng/mL GM-CSF (ThermoFisher #300-03), was used, with media being changed every other day. iMicroglia were considered mature after 10 days in maturation media and used for assays within 5 ± 4 days post maturation.

### Dopaminergic-like Neuron Differentiation

For production of neuronal debris as a positive inducer of microglia activation, we differentiated the Lund human mesencephalic (LUHMES) cell line into dopaminergic-like neurons previously published^107^, with modifications. In brief, LUHMES cells were cultured in DMEM/F12 (Gibco #11320-033) with N2 (Gibco #17502048), 20 mM L-Glutamine (Invitrogen #25030-081), and 40 ng/mL FGF (Invitrogen #PHG0021) in 75 cm^2^ flask coated with 50 ug/mL Poly-L-Ornithine Hydrobromide (PLO) (Sigma #P3655-100MG) and 1 μg/mL Human Plasma Fibronectin (Gibco #33016015). Once cells reached 80-90% confluency, cells were lifted with a 1:1 dilution of TrypLE^TM^ Express (Gibco #12604-013) and Phosphate Buffered Saline (PBS, Gibco #10010-023) and replated 2.6 x 10^7^ cells into a 175 cm^2^ flask with the same PLO-Fibronectin coating with LUHMES differentiation media of DMEM/F12 (Gibco #11320-033) with N2 (Gibco #17502048), 20 mM L-Glutamine (Invitrogen #25030-081), 1 mM Dibutyryl cAMP (Sigma #D0627), 1 μg/mL doxycycline (Sigma-Aldrich #D3447), and 2 ng/mL GDNF (Sigma #SRP3309-10UG). After 48-hours, cells were replated into a new 175 cm^2^ flask in fresh differentiation media at a density of 26 x 10^6^ cells. Neuronal differentiation was considered complete after an additional three days, consistent with previous publications^108–110^. Successful differentiation was visually confirmed by morphological changes, including attachment to the dish and growth of axon-like processes. Differentiated cells were then used within 3 ± 2 days post maturation.

### Production of Damage-Associated Molecular Patterns (DAMPs) from Neurons

To produce Damage-Associated Molecular Patterns (DAMPs) from neurons, we used an Apoptotic/Neuritic Neuron induction protocol previously described^30,51^. In brief, differentiated dopaminergic neurons from the LUHMES cell line were exposed to UV-B at 306 nm for 15 minutes (Boekel Scientific #234100). After 24 hours, cells were floating or detached by gently pipetting around the well with PBS. Cells were centrifuged at 1000 g for 5 minutes and resuspended in PBS for immediate use.

### Exposure of iMicroglia to biological and chemical perturbations

Mature iMicroglia were replated into 6-well dishes at an average density of 400,000 per well (100,000 – 700,000 variable by line) with fresh media. Twenty-four hours after replating, cells were exposed to chemical/drugs (resuspended in DMSO), DAMPs (resuspended in PBS), or left in microglia maturation media as a control. Exposures of HDAC inhibitors (Vorinostat, Ambeed #122285; Entinostat, Ambeed #A234507) were determined by the literature^57^, 0.1 μM and 1 μM each. DAMPs produced from LUHMES were counted using a 5 ug/mL concentration of Acridine Orange/Propidium Iodide Stain (Logos #F23001) in ultra-low fluorescence counting slides (PhotonSlide #L12005), and added at 0.5X, 1X, and 2X cells per cm^2^ (equal to 168,000, 336,000, and 672,000 cells per well in a 6-well plate) previously reported to induce DAM-like expression^51^ (35,000 cells per cm^2^). Sub-toxic exposures of Inteferon gamma (Thermo Fisher #300-02 - 2.5 ng/mL, 12.5 ng/mL, and 25 ng/mL), PIKfyve inhibitors (Apilimod, Ambeed #A149227 - 0.01 μM, 0.1 μM, 1μM; Vacuolin, Ambeed #A1359469 - 0.1 μM, 1μM, 10 μM), and BTK inhibitors (Ibrutinib, Ambeed # A150396 - 0.01 μM, 0.1 μM, 1μM; Spebrutinib, Ambeed #A537377 - 0.1 μM, 1μM, 10 μM) were determined by CellTitre-Glo® 2.0 Cell Viability Assay (Promega, G9241; Data Not Shown). After 24 hours, cells were processed for transcriptomic profiling.

### RNA isolation, library preparation, and mRNA sequencing

RNA from exposed cells and controls was isolated using the Qiagen RNEasy Plus Mini kit (Qiagen, #74134). After aspirating media, cells were lysed in the well per manufacturer protocol. Genomic DNA cleanup and RNA isolation were performed per recommended protocol without modification. Isolated RNA was quantified via spectrophotometer (NanoDrop ND-2000) and transferred to the National Heart, Lung and Blood Institute’s (NHLBI) DNA Sequencing and Genomics core facility (Bethesda, MD).

Library preparation for mRNA sequencing was performed using Illumina Stranded mRNA Prep Kit (Illumina #20040534) per manufacturer protocol. Quality control for sample distribution was performed with MiSeq (MiSeq Reagent Nano Kit v2 (300-cycles); Illumina MS-103-1001) and subsequently sequenced with NovaSeq X Series 1.5B Reagent Kit (200 cycles): (Illumina #20104704) set for a paired-end 100 bp run (100:10:100:10 cycle run) to a mean sequencing depth of ∼30M reads per sample (± 6.5, mean ± standard deviation).

### Bulk RNA-seq data preprocessing and quality control

Fastqs were aligned and quantified using nf-core/rnaseq (v3.19.0)^111^ of the nf-core collection of workflows^112^, utilising reproducible software environments from the Bioconda^113^ and Biocontainers ^114^ projects. The pipeline was executed with Nextflow (v25.04.2)^115^. Briefly, reads were trimmed using trimgalore (v0.6.10) (https://github.com/FelixKrueger/TrimGalore), aligned to the same reference genome used for snRNA-seq preprocessing (CellRanger reference 2024-A, GRCh38v110) using STAR (v2.6.1d)^116^ and quantified using Salmon (v1.10.3)^117^.

### Differential expression analysis of treated iMicroglia

Prior to differential expression analysis, low-expression genes were filtered. Briefly, for a given batch of data, counts were TMM-normalized using the *edgeR* package, and genes were retained if they were expressed at a CPM > 1 in at least three samples of any treatment group (including controls). Differential expression was then performed using *dream*^52^, a differential expression framework for bulk RNAseq analysis built off the limma-voom pipeline^118^ modified to accommodate random effect variables. In brief, each [treatment x dose] (e.g., Entinostat_1uM) was tested against batch-specific control samples with biological replicates (n = 5 PPMI lines, independent differentiations). The formula for differential expression included a random effect to account for line variability in plating densities and cell line proliferation (formula: GEx ∼ treatment_dose + (1 | PPMI_line)). Adjusted p-values were used from each independent comparison, and genes were considered significantly differentially expressed with an adjusted p-value < 0.05.

### Comparison to *in vivo* human signatures

As described above (See **“Characterization of cell subtypes”**), gene set enrichment analysis was performed for a signed, quantitative approach to describing the presence of gene signatures using *fgsea*^84^. Results from differential expression in bulk RNA from treated *in vitro* studies were analyzed against the KEGG “Apoptosis” pathway (KEGG hsa04210) and the hnDAM signature as defined in (**Figure 4**).

## Resource Availability

### Lead Contact

Further information and requests for resources and reagents should be directed to and will be fulfilled by the lead contact, Mark R. Cookson.

### Materials Availability

No physical material or resources were produced by this study.

### Data and Code Availability

No human postmortem data was collected from this study. Transcriptomic data were accessed where publicly available or under specific data use agreements (**Table S38**). Since the datasets used in this study come with highly variable data use and privacy requirements, no complete sample- or individual-level processed data can be made available. However, code necessary to replicate all analyses is publicly available on GitHub (https://github.com/neurogenetics/Glia_across_NDDs).Summary level information may be found in the Supplementary Information associated with this manuscript or a stable release found on Zenodo (https://doi.org/10.5281/zenodo.17203360). Data from sequencing of cell lines differentiated to iPSC microglia will be made available on the Parkinson’s Progression Markers Initiative (PPMI) database (www.ppmi-info.org/data) under PPMI data use agreements upon acceptance for publication.

## Acknowledgements

We would like to thank Xylena Reed, Changyoun Kim, and Elise Marsan for their feedback and thoughtful discussion concerning transcriptomic analyses of glial cells. We would also like to thank David Gerhold, Zhibin Tong, and Maria Garcia Aguilar for their guidance on differentiating dopaminergic neurons from the LUHMES cell line. This research was supported by the Intramural Research Program of the National Institute on Aging (Z01AG000931, Z01AG000953 to M.R.C.), funding from the Michael J. Fox Foundation (MJFF-025642), and data used in the preparation of this article were obtained from the Parkinson’s Progression Markers Initiative (PPMI) database (www.ppmi-info.org/data; for full acknowledgment see **Supp Data**), RRID:SCR_006431. For up-to-date information on the study, visit http://www.ppmi-info.org. This work utilized the computational resources of the NIH HPC Biowulf cluster (https://hpc.nih.gov). Schematics within figures and graphical abstract created with <u>BioRender.com</u>.

We would also like to thank the patients, patient families, and physicians who contributed to the resources used in this manuscript. Information regarding publicly available datasets used without data use agreements from the Gene Expression Omnibus or Human Cell Atlas are available in the Supplementary Information (**Table S38**). We would also like to acknowledge the work on several consortia-level programs: Accelerating Medicine Partnership® (AMP®) Parkinson’s Disease (AMP PD), Alzheimer’s Disease (AD) Knowledge Portal, the Religious Orders Study and Rush Memory and Aging Project (ROSMAP), and Seattle-AD (SEA-AD). Full acknowledgement of respective studies, brain banks, and data use agreements available in the Supplementary Information (**Supp Data**).

This research was supported by the Intramural Research Program of the National Institutes of Health (NIH). The contributions of the NIH authors were made as part of their official duties as NIH federal employees, are in compliance with agency policy requirements, and are considered Works of the United States Government. However, the findings and conclusions presented in this paper are those of the authors and do not necessarily reflect the views of the NIH or the U.S. Department of Health and Human Services.

## Ethics and Generative AI Use Statement

Large language models (LLMs) were employed for optimization or annotation of code, under the current best practice guidance of the NIH (https://osp.od.nih.gov/policies/artificial-intelligence/). We, the authors, affirm that no LLMs were used in the study design, drafting of any text, or analysis of individual-level or de-identified human data.

## Author Contributions

L.H-P., D.J.A., and M.R.C. conceptualized the study and planned experiments. Investigation and formal analysis was performed by L.H-P., D.J.A., A.M., J.B., H.M.B., A.R.D., and J.R.G.. Funding of this study was obtained by M.R.C. and the first draft of the manuscript was written by L.H-P. and D.J.A.. All authors reviewed and edited the manuscript. M.R.C. and A.R.D. supervised the work.

## Supplementary Information

Supplementary figures include comparisons of glial taxonomy to published signatures and astrocyte subclustering (Figs. S1-2). Initial characterizations and disease associations are supported by proportion analysis, meta-analysis with mashr, cluster markers, and detailed cell counts (Fig. S3; Tables S1-13). An overview of pseudobulked differential expression, covariate selection, meta-analysis with mashr, and pathway analysis are available online (Figs S4-8; Tables 14-28). To define the hnDAM signature information includes latent variable discovery, pathway analysis, transcription factor analysis, validation in an *in vivo* replication cohort, and transcriptional programming discovery (Figs S9-10, Tables S29-40). Results pertaining to *in vitro* perturbation of iPSC-derived microglia (Tables S41-45), analysis of an *in vitro* replication cohort (Tables S48-50), and comparison to lecanemab-induced microglia activation (Tables S51-52) are also available online. A stable release of the supplementary information associated with this manuscript can be found on Zenodo (10.5281/zenodo.17203361).

## Supplemental Data

### AMP-PD

Data used in the preparation of this article were obtained from the Accelerating Medicine Partnership® (AMP®) Parkinson’s Disease (AMP PD) Knowledge Platform. For up-to-date information on the study, visit https://www.amp-pd.org.

The AMP® PD program is a public-private partnership managed by the Foundation for the National Institutes of Health and funded by the National Institute of Neurological Disorders and Stroke (NINDS) in partnership with the Aligning Science Across Parkinson’s (ASAP) initiative; Celgene Corporation, a subsidiary of Bristol-Myers Squibb Company; GlaxoSmithKline plc (GSK); The Michael J. Fox Foundation for Parkinson’s Research ; Pfizer Inc.; AbbVie Inc.; Sanofi US Services Inc.; and Verily Life Sciences. ACCELERATING MEDICINES PARTNERSHIP and AMP are registered service marks of the U.S. Department of Health and Human Services.

### AD Knowledge Portal

The results published here are in whole or in part based on data obtained from the AD Knowledge Portal (https://adknowledgeportal.org/).

### The MIT ROSMAP Single-Nucleus Multiomics Study

Study data were generated from postmortem brain tissue provided by the Religious Orders Study and Rush Memory and Aging Project (ROSMAP) cohort at Rush Alzheimer’s Disease Center, Rush University Medical Center, Chicago. This work was supported in part by the Cure Alzheimer’s Fund, NIH grants AG058002, AG062377, NS110453, NS115064, AG062335, AG074003, NS127187, MH119509, HG008155 (M.K.), RF1AG062377, RF1 AG054321, RO1 AG054012 (L.-H.T.) and the NIH training grant GM087237 (to C.A.B.). ROSMAP is supported by P30AG10161, P30AG72975, R01AG15819, R01AG17917. U01AG46152, U01AG61356.

### ROSMAP

Study data were generated from postmortem brain tissue provided by the Religious Orders Study and Rush Memory and Aging Project (ROSMAP) cohort at Rush Alzheimer’s Disease Center, Rush University Medical Center, Chicago. This work was funded by NIH grants U01AG061356 (De Jager/Bennett), RF1AG057473 (De Jager/Bennett), and U01AG046152 (De Jager/Bennett) as part of the AMP-AD consortium, as well as NIH grants R01AG066831 (Menon) and U01AG072572 (De Jager/St George-Hyslop).

### Seattle-AD

Study data were generated from postmortem brain tissue obtained from the University of Washington BioRepository and Integrated Neuropathology (BRaIN) laboratory and Precision Neuropathology Core, which is supported by the NIH grants for the UW Alzheimer’s Disease Research Center (P50AG005136 and P30AG066509) and the Adult Changes in Thought Study (U01AG006781 and U19AG066567). This study is supported by NIA grant U19AG060909.

### PPMI (Parkinson’s Progression Markers Initiative) Biospecimen Use Agreement

Data biospecimens used in the analyses presented in this article were obtained from the Parkinson’s Progression Markers Initiative (PPMI) (www.ppmi-info.org/access-dataspecimens/download-data). As such, the investigators within PPMI contributed to the design and implementation of PPMI and/or provided data and collected biospecimens, but did not participate in the analysis or writing of this report. For up-to- date information on the study, visit www.ppmi-info.org. PPMI – a public-private partnership – is funded by the Michael J. Fox Foundation for Parkinson’s Research and funding partners, including 4D Pharma, Abbvie, AcureX, Allergan, Amathus Therapeutics, Aligning Science Across Parkinson’s, AskBio, Avid Radiopharmaceuticals, BIAL, BioArctic, Biogen, Biohaven, BioLegend, BlueRock Therapeutics, Bristol-Myers Squibb, Calico Labs, Capsida Biotherapeutics, Celgene, Cerevel Therapeutics, Coave Therapeutics, DaCapo Brainscience, Denali, Edmond J. Safra Foundation, Eli Lilly, Gain Therapeutics, GE HealthCare, Genentech, GSK, Golub Capital, Handl Therapeutics, Insitro, Jazz Pharmaceuticals, Johnson & Johnson Innovative Medicine, Lundbeck, Merck, Meso Scale Discovery, Mission Therapeutics, Neurocrine Biosciences, Neuron23, Neuropore, Pfizer, Piramal, Prevail Therapeutics, Roche, Sanofi, Servier, Sun Pharma Advanced Research Company, Takeda, Teva, UCB, Vanqua Bio, Verily, Voyager Therapeutics, the Weston Family Foundation and Yumanity Therapeutics.

---

**Fig. S1.**
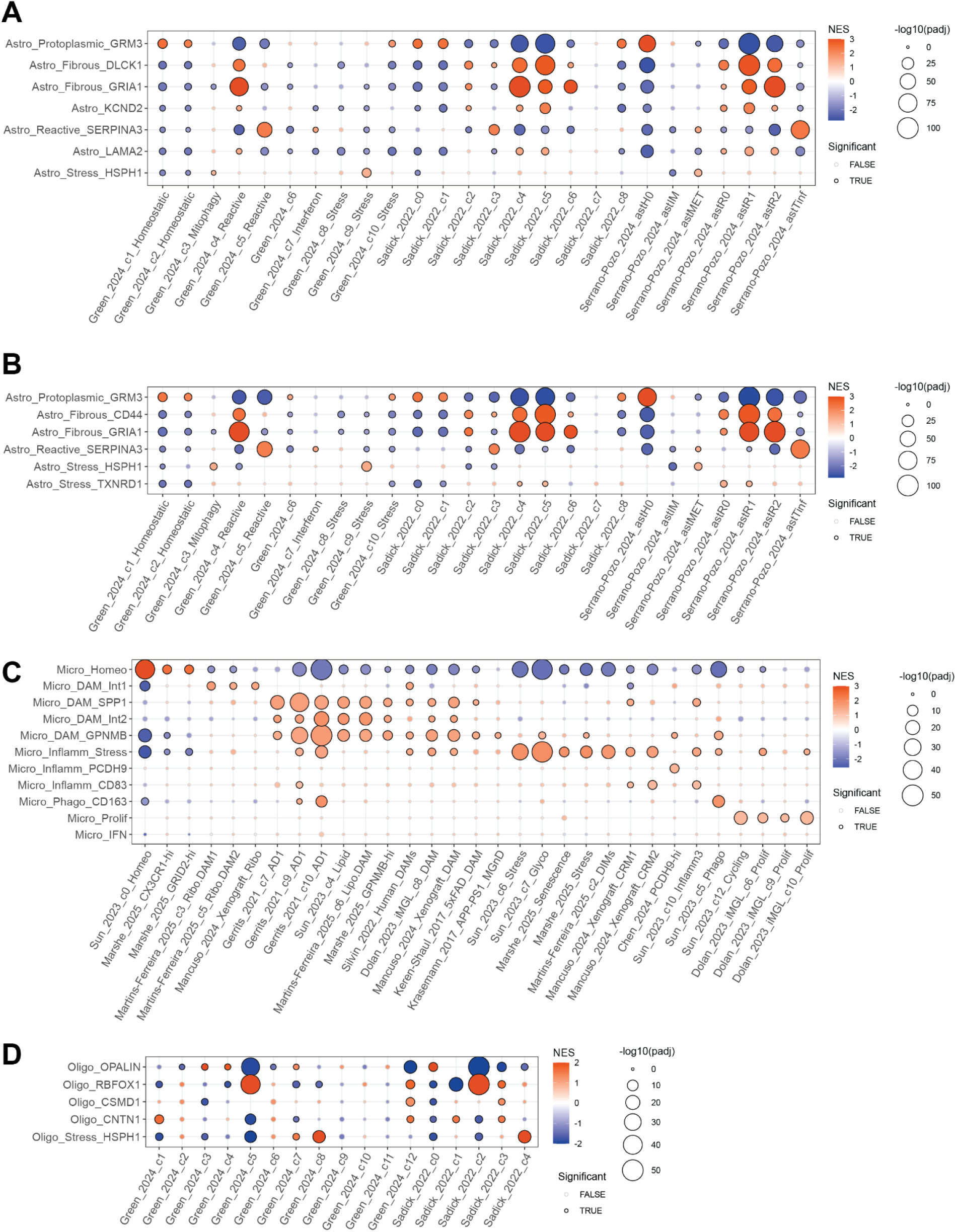
Cross-NDD glial substate diversity reflected in published human and mouse signatures. GSEA-style enrichment performed in *fgsea* against published disease-associated glial subtypes (see **“Characterization of cell subtypes”**). (**A**) Total and (**B**) cortical astrocyte annotations segregated mainly by fibrous (*GFAP*+) and protoplasmic (*GFAP*-) recapitulate 26 known astrocytic substates from three human studies. (**C**) Unified naming strategy captures the transcriptional state of 31 microglial substates across seven human and two mouse studies. (**D**) *De novo* oligodendrocyte clusters recapitulates limited known heterogeneity of 17 oligodendrocytic sub-states from two human studies.

**Fig. S2.**
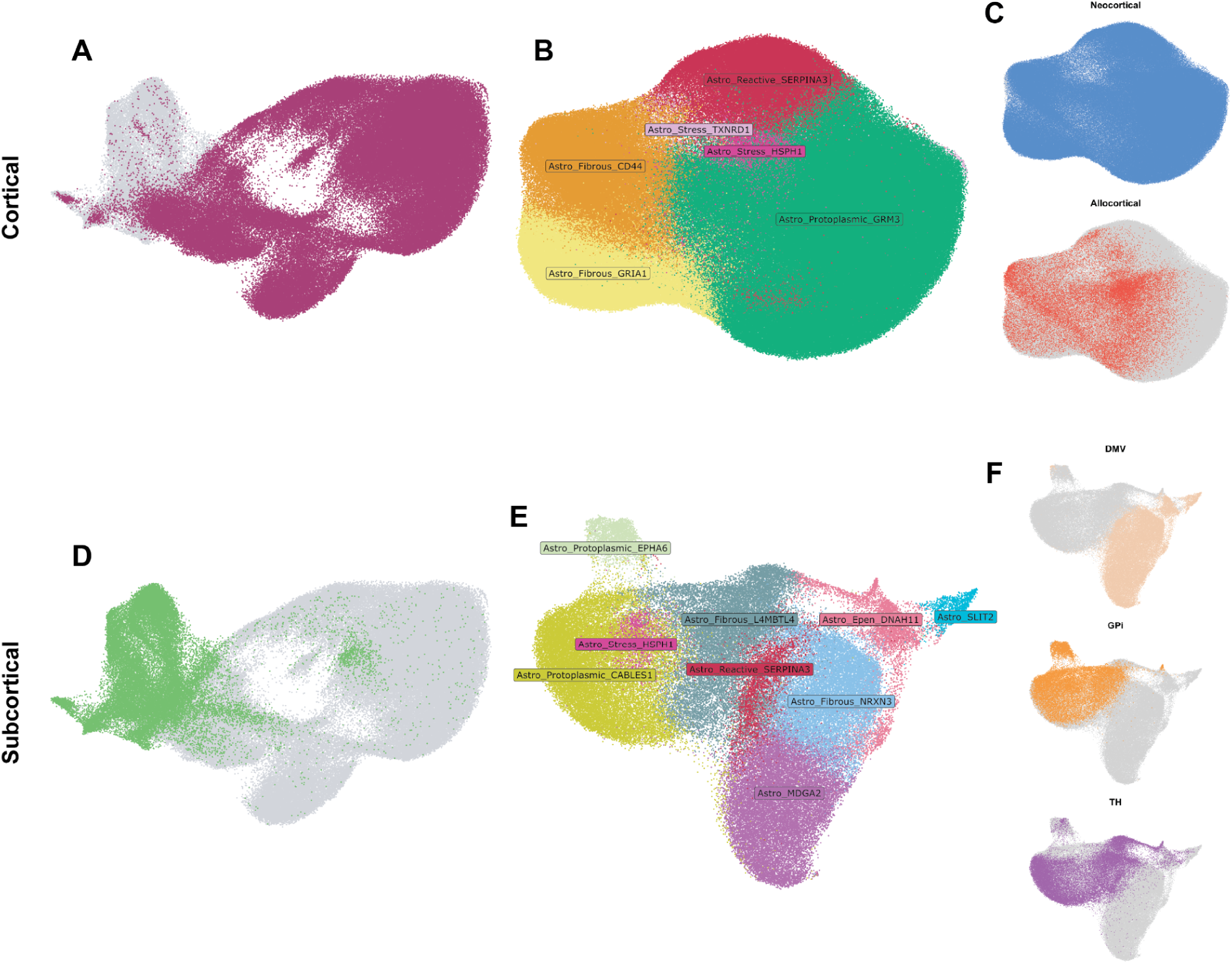
Astrocyte transcriptional state driven by regional heterogeneity. **(A)** Cortical astrocytes (HIP, EC, AnG, M1, FC/PFC, OC/V1, TC/MTG) from initial clustering (Fig. 1A). (**B**) Successive subclustering reveals 6 clusters at a resolution of 0.15. (**C**) Allocortical (HIP/EC) and neocortical regions (AnG, M1, FC/PFC, OC/V1, TC/MTG) display minimal sub-regional heterogeneity. (**D**) Subcortical astrocytes (DMV, GPi, TH) from initial clustering (Fig. 1A). (**E**) Successive sub-clustering reveals 9 clusters at a resolution of 0.2. (**F**) Substantial sub-regional heterogeneity apparent across 3 subcortical regions.

**Fig. S3.**
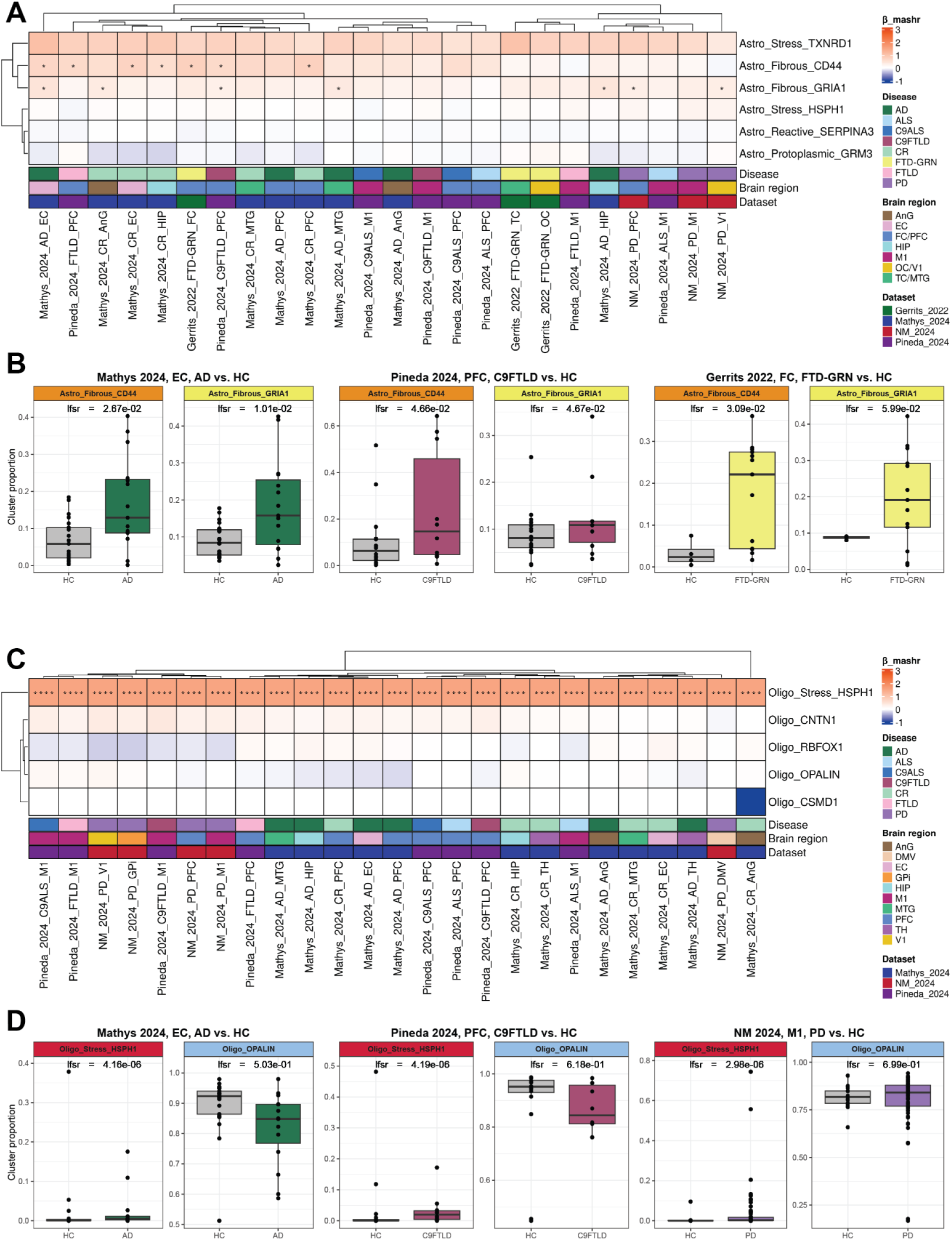
Astrocyte and oligodendrocyte subpopulations lack strong cross-NDD associations. Differential abundance of subtype populations was tested using a GLM covaried by sex and age and β and p-values were refined using *mashr*^90^. (**A**) Astrocytic differential abundance reveals 2 disease-associated subtypes across NDD and regions (* LFSR *< 0.05*) (**B**) Per sample proportion changes of DAA-like subtypes in AD, C9TLD, FTD-GRN (CITE). (**C**) Oligodendrocytic differential abundance reveals 1 disease-associated subtypes across NDD and regions (**** LFSR *< 0.0001*) (**D**) Per sample proportion changes of DAO-like subtypes in AD, C9TLD, PD (CITE). See **Table S13** for summary statistics.

**Fig. S4.**
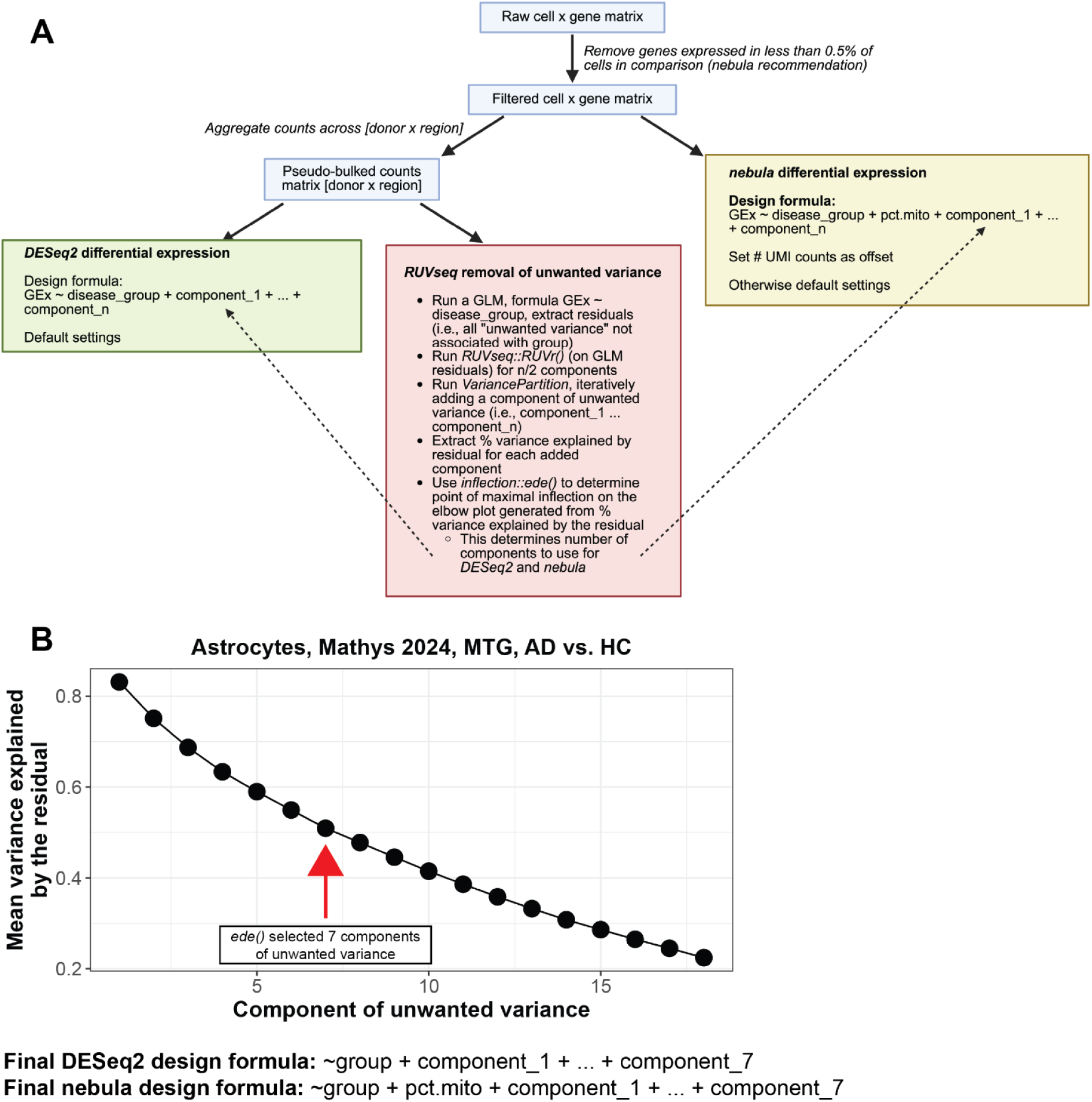
Overview of differential expression workflow. (**A**) Feature and sample filtering strategies for pseudo-bulk and cell-level differential expression. Latent factors to be included as covariates were selected automatically using *RUVseq* (See **“Differential gene expression analysis”**). (**B**) Example of automated hyperparameter selection to define components of unwanted variance.

**Fig. S5.**
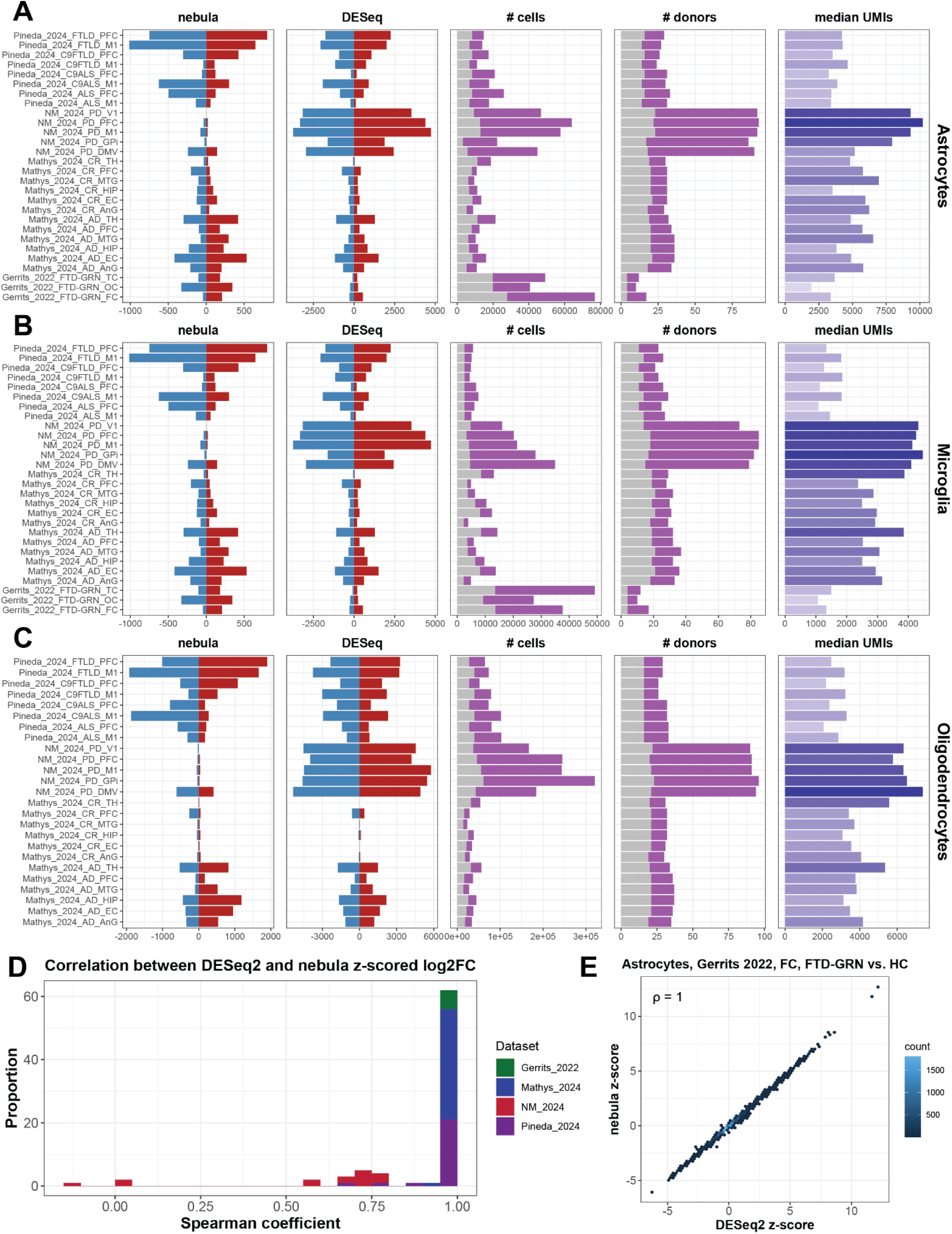
Summary of pseudobulk and cell-level differential expression results. (**A**-**C**) Number of differentially expressed genes at the summarized, number of cells past QC filters, number of donors contributing cells, and median UMIs per condition [disease x region] across respective glial types. (**D**) A histogram of spearman correlation of *DESeq2* and *nebula* results per condition [disease x region]. (**E**) Correlation of z-scored log2FC for select comparison reveals correlation coefficient equal to one (Spearman ⍴ = 1).

**Fig. S6.**
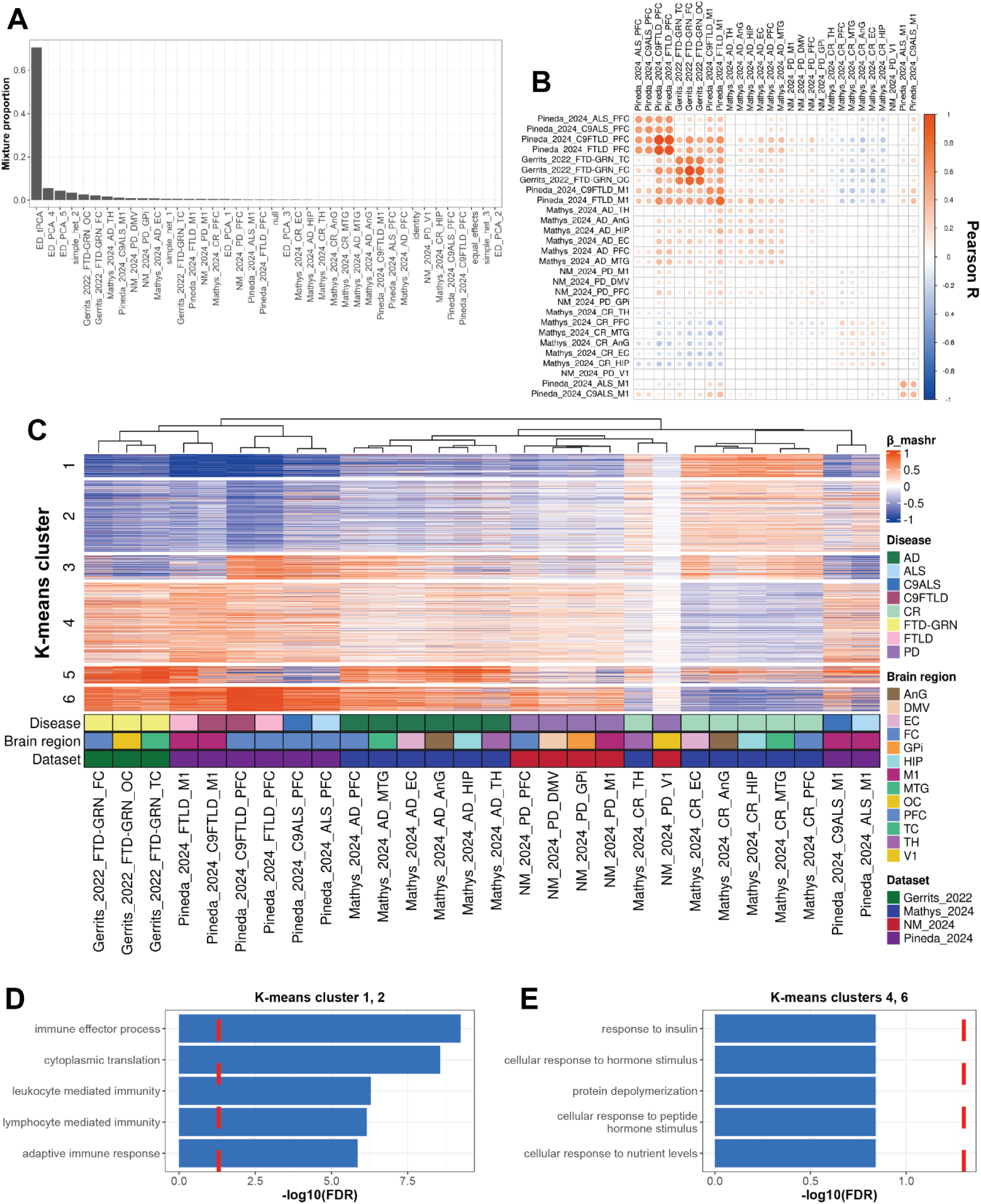
Cross-disease meta-analysis of microglial gene expression reveals unique association with cognitive resilience. (**A**) Mixture proportions of different types of covariance matrices generated from *mashr* analysis of microglia reveals data-driven model (ED_tPCA) explains the most sharing of cell-level differential expression profiles (computed by *nebula*). (**B**) Covariance matrix demonstrates patterns of shared variance explained by ED_tPCA. (**C**) K-means clustering (k = 6) of genes with sharing most explained by ED_tPCA (n = 2,500). GO terms computed in *ClusterProfiler* reveal immune-processes in the union of clusters one and two (**D**) and no significantly enriched pathways in the union of clusters four and six (**E**).

**Fig. S7.**
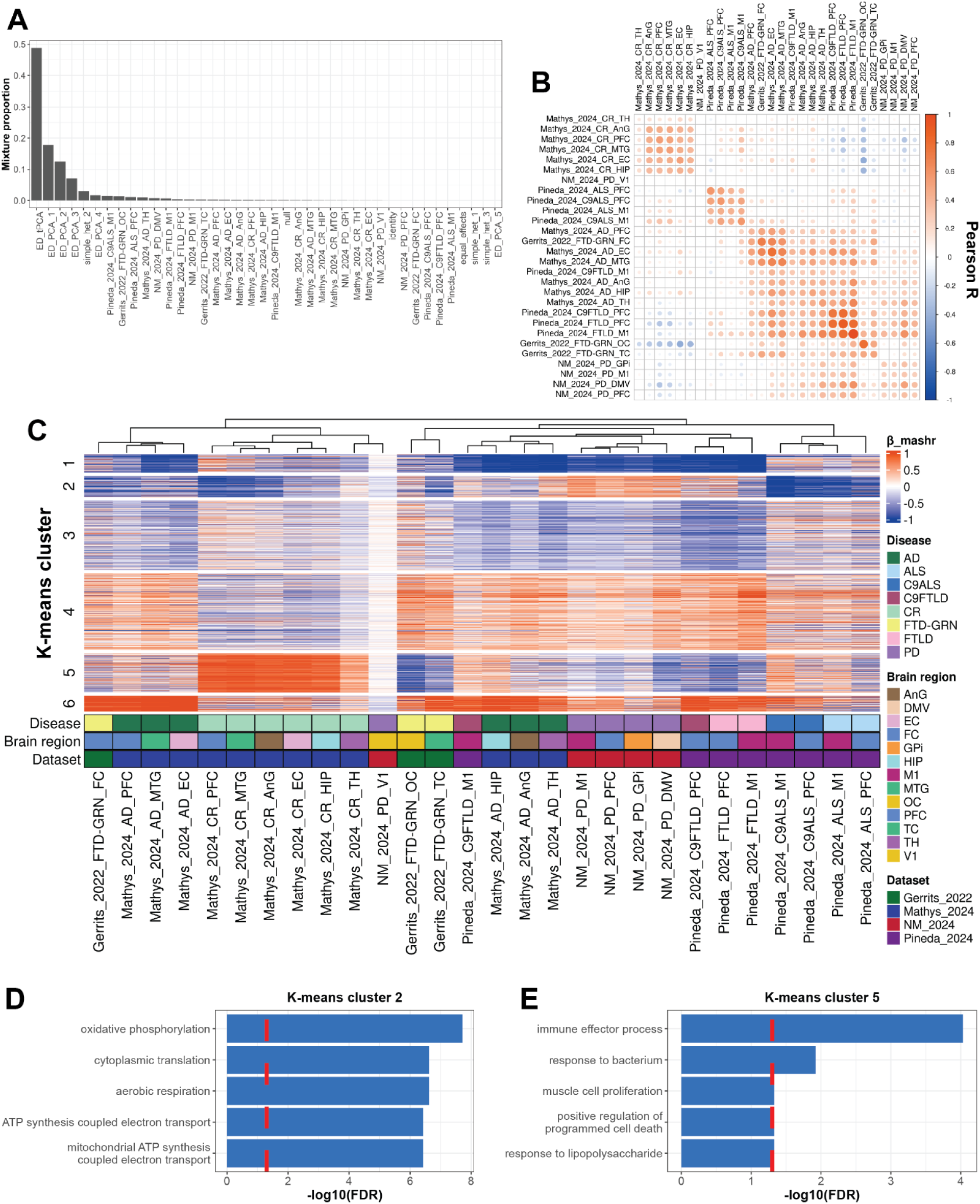
Cross-disease meta-analysis of astrocyte gene expression reveals associations with ALS and cognitive resilience. (**A**) Mixture proportions of different types of covariance matrices generated from *mashr* analysis of astrocytes reveals data-driven model (ED_tPCA) explains the most sharing of cell-level differential expression profiles (computed by *nebula*). (**B**) Covariance matrix demonstrates patterns of shared variance explained by ED_tPCA. (**C**) K-means clustering (k = 6) of genes with sharing most explained by ED_tPCA (n = 3,502). GO terms computed in *ClusterProfiler* reveal aerobic respiration in cluster two (**D**) and immune-related pathways in cluster five (**E**).

**Fig. S8.**
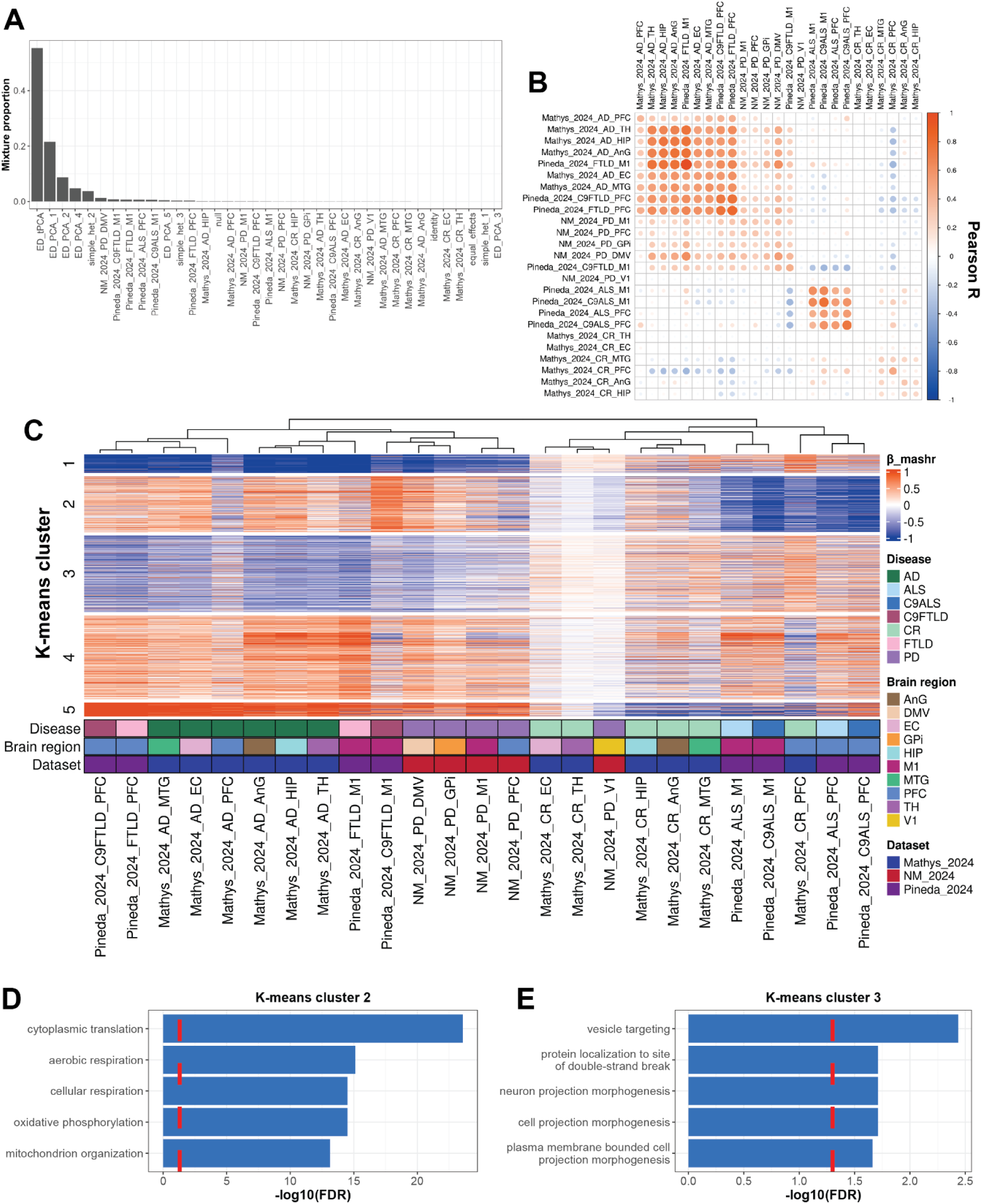
Cross-disease meta-analysis of oligodendrocyte gene expression reveals associations with ALS. (**A**) Mixture proportions of different types of covariance matrices generated from *mashr* analysis of oligodendrocytes reveals data-driven model (ED_tPCA) explains the most sharing of cell-level differential expression profiles (computed by *nebula*). (**B**) Covariance matrix demonstrates patterns of shared variance explained by ED_tPCA. (**C**) K-means clustering (k = 5) of genes with sharing most explained by ED_tPCA (n = 1,238). GO terms computed in *ClusterProfiler* reveal aerobic respiration in cluster two (**D**) and membrane trafficking-related pathways in cluster three (**E**).

**Fig. S9.**
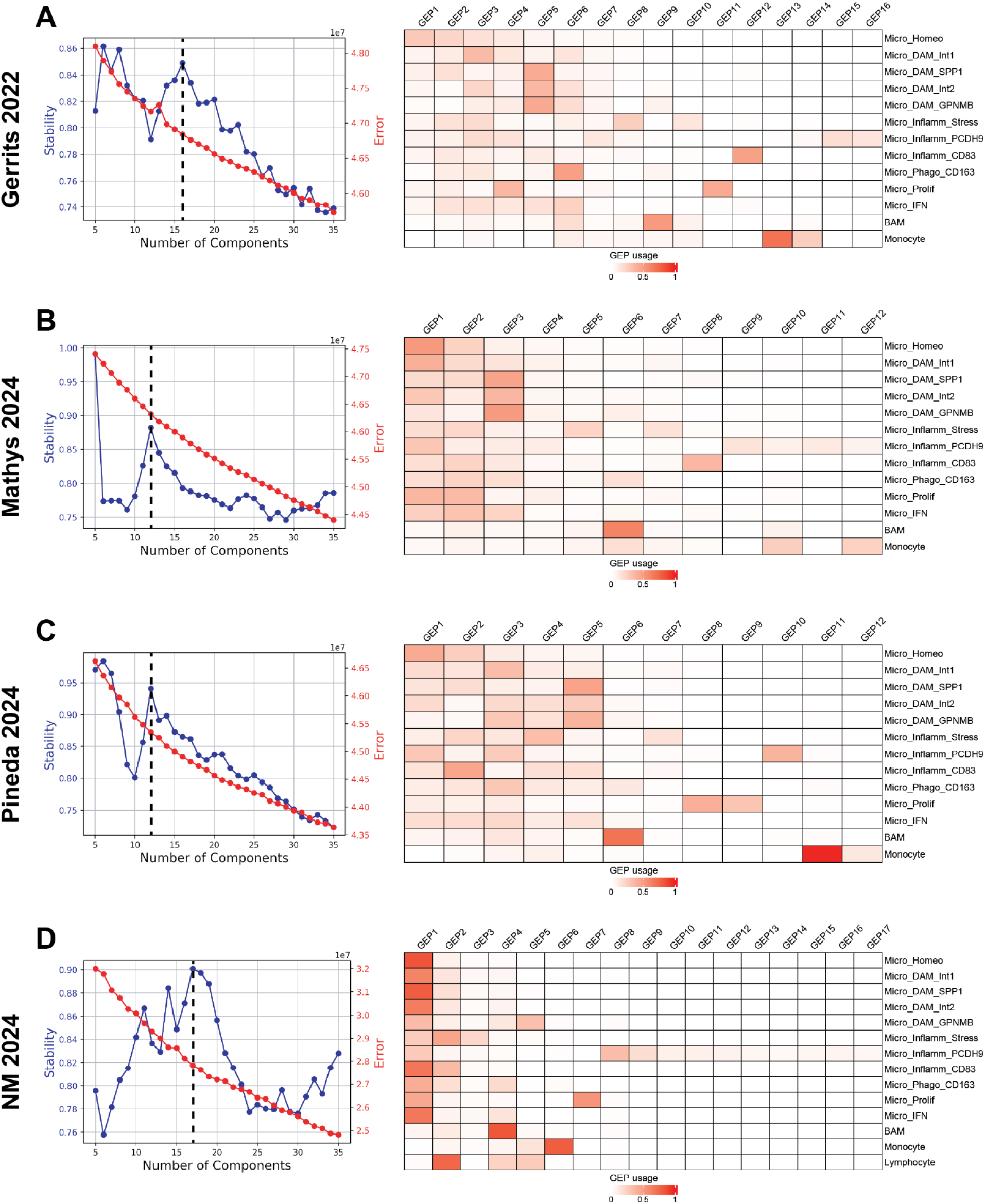
Latent factorization of microglial gene expression programs. The definition of the hnDAM module was contingent on the discovery of latent factors shared across disease-associated subtypes. Consensus non-negative matrix factorization allowed for the discovery of factors within each study of the discovery cohort (See **Methods**). The number of gene expression programs (GEPs) were determined: (**A**) 16 GEPs for Gerrits_2022, (**B**) 12 GEPs for Mathys_2024, (**C**) 12 GEPs for Pineda_2024, (**D**) 17 GEPs for NM_2024. Heatmaps of scaled GEP usage across microglia substates represented as heatmaps.

**Fig. S10.**
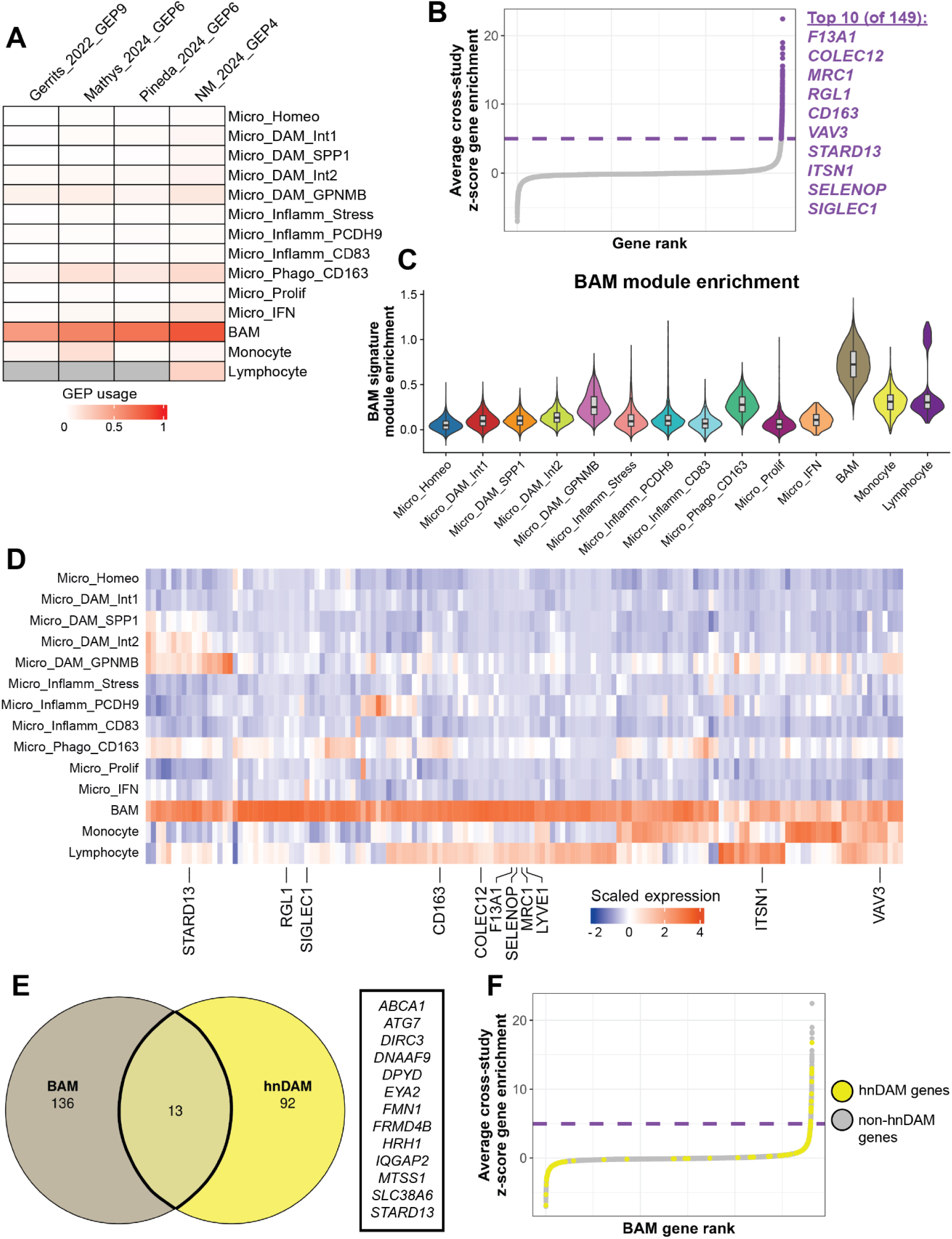
Cross-study identification of a border-associated macrophage gene expression program. Microglial gene expression programs were defined using *cNMF*^100^ (see “**Microglia signature discovery via latent factorization”**). (**A**) GEPs uniquely enriched in border-associated macrophages were apparent from all four discovery studies. (**B**) Genes were prioritized from Gerrits_2022_GEP9, Mathys_2024_GEP6, Pineda_2024_GEP6, and NM_2024_GEP4 by z-scoring eigenvalues within respective GEPs. Z-scores for each gene were averaged across study to define the BAM signature (149 genes, average z-score > 5). (**C-D**) Expression of BAM genes across microglial subtypes shows enrichment in one BAM cluster. A module score was assigned to each cell using the *AddModuleScore()* function (**C**), or expression was computed using the *AverageExpression()* function in *Seurat*, and z-scored across subtypes (**D**). (**E**) Comparison of BAM signature to hnDAM (Fig. 3) highlights discrete gene membership. (**F**) hnDAM genes are neither reversed (negative value), nor similarly enriched (positive value) when ranked by cross-study z-score of BAM signature.

**Fig S11.**
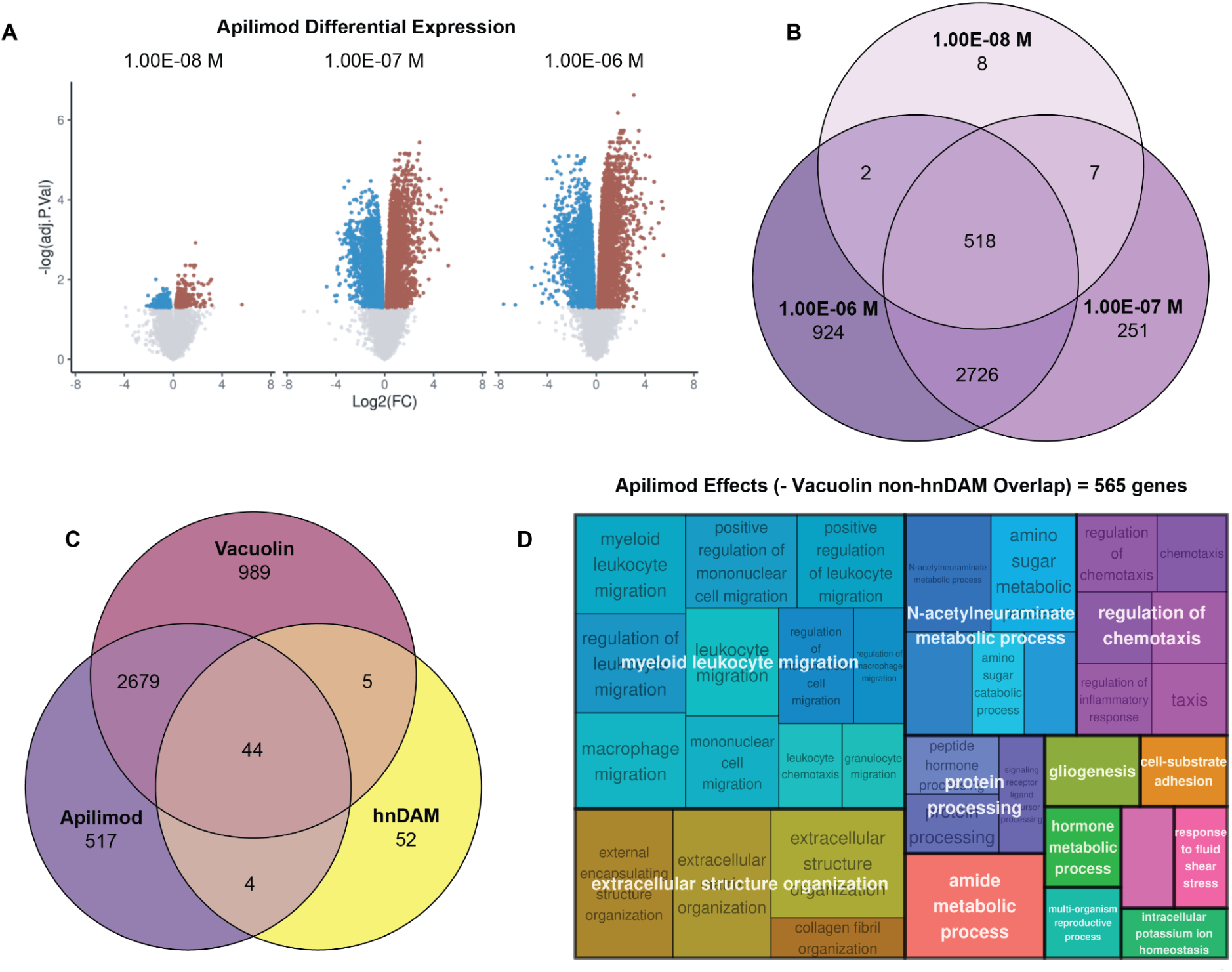
Apilimod induces cell motility and metabolism pathways in iPSC-derived microglia. Differential expression performed after 24 hour treatment of Apilimod in dream (∼ treatment_dose) available in **Table S44**. (**A**) Number of down-regulated (blue; FDR < 0.05 and logFC < -0.5) and up-regulated genes (red; FRD < 0.05 and logFC > 0.5) increase with treatment dose. (**B**) Strong overlapping effects observed with 1 μM (1E-6 M) and 100 nM (1E-7 M) treatment including 48 of 105 hnDAM genes. (**C**) Apilimod and PIKfyve inhibitor analog Vacuolin have shared and distinct effects, but jointly regulate 44 of 105 hnDAM genes. Apilimod and vacuolin effects summarized as common DEGs between 1 μM and 100 nM treatments for their respectively. (**D**) GO enrichment of distinct apilimod-specific effects are significantly enriched for cell motility and metabolism pathways (BH < 0.05) reduced using *rrvgo*. Full enrichment results available in **Tables S46-47**.

**Fig S12.**
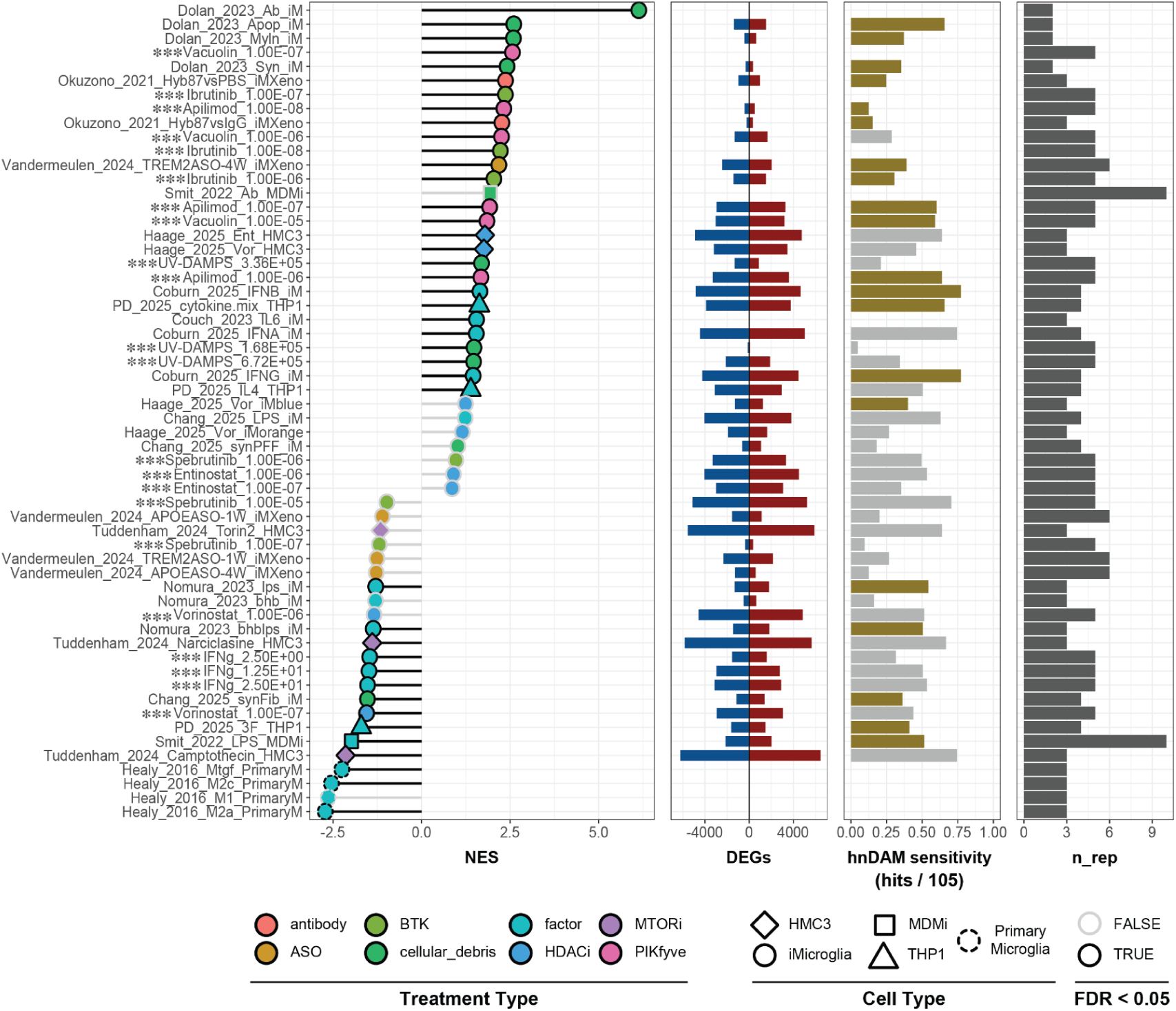
*In vitro* replication cohort reveals hnDAM induction across myeloid cell models. hnDAM signal was queried in independently collected sample series across NDDs (12 studies: **Table S48**) using GSEA-based scoring (See **“Validation and replication of hnDAM signature”**). Differential expression results summarized for effects FDR < 0.05 and absolute value of logFC > 0.5 (**Tables S49**). Hypermetric test reveals significant enrichment to hnDAM signature (gold = FDR < 0.05) plotted with sensitivity (percent hnDAM genes listed as DEG) and sample size reported for each study (n_rep). P-value reported from *fgsea* were Benjamini-Hochberg FDR-corrected across all 34 comparisons in the replication cohorts and 22 comparisons from the present study (*** prefix). Enrichment statistics available in **Table S50**.

**Fig S13.**
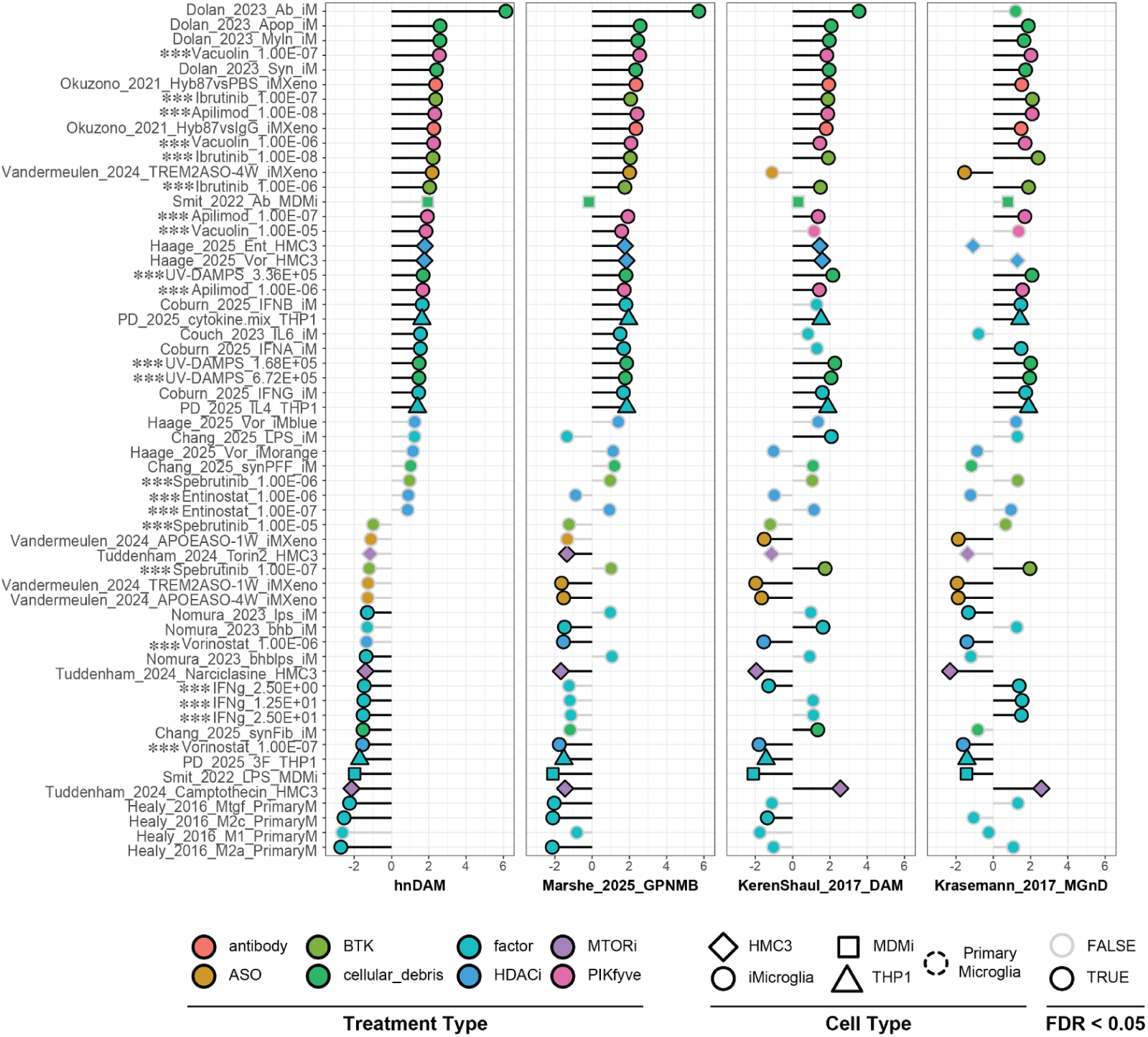
Human and mouse DAM signatures are induced by distinct treatment paradigms. hnDAM signal was queried in independently collected sample series across NDDs (12 studies: **Table S48**) using GSEA-based scoring (See **“Validation and replication of hnDAM signature”**). Pathway analysis faceted by pathway of interest (hnDAM, Marshe_2025_GPNMB, Keren-Shaul_2017_DAM, and Krasemann_2017_MGnD). Enrichment statistics available in **Table S50**.

**Fig S14.**
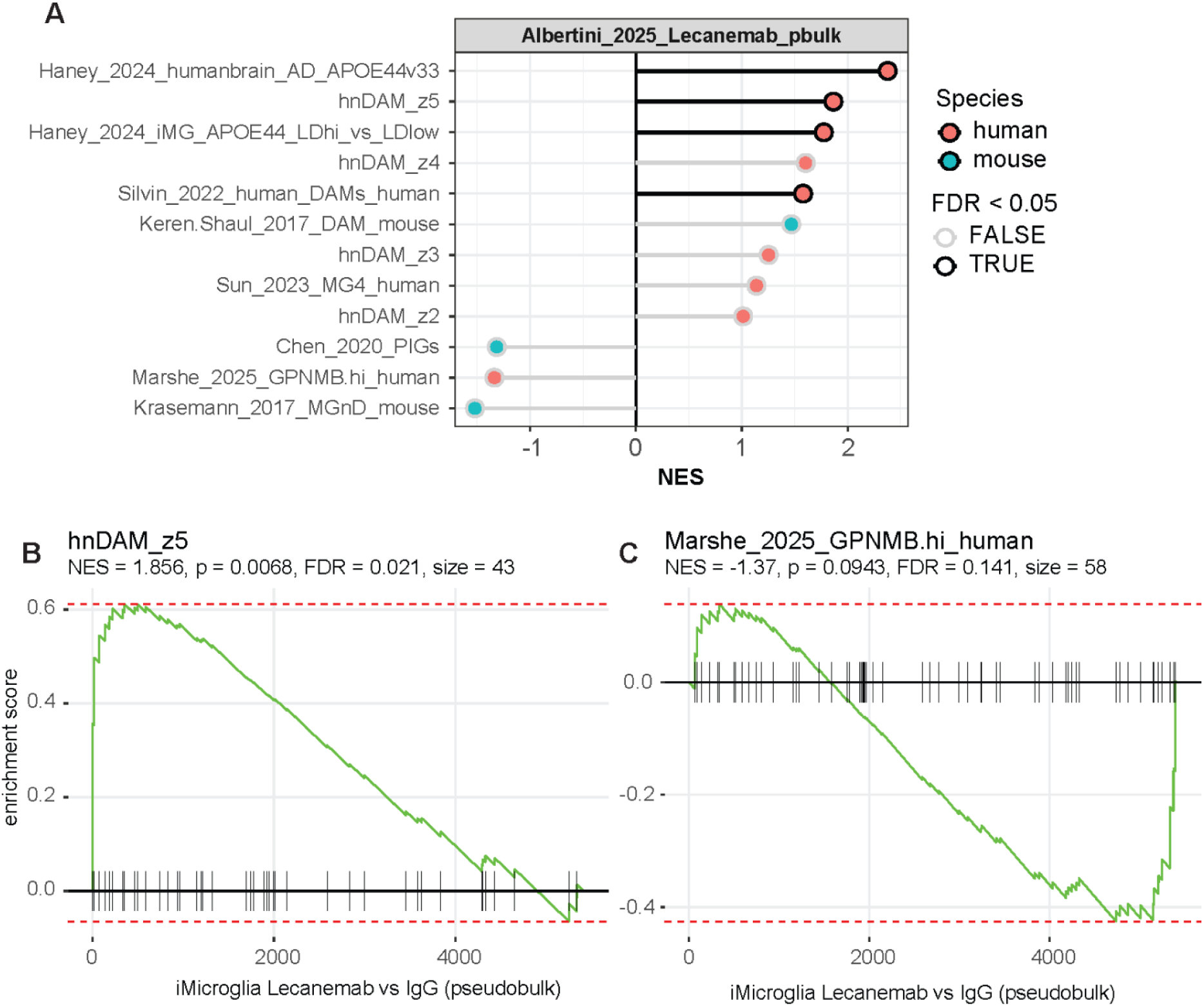
Lecanemab-induced microglia shifts reflect distinct human DAM signatures. hnDAM signal was queried publicly available pseudobulk differential expression results comparing Lecanemab treated microglia versus IgG control after xenograft into an amyloid mouse model ^66^. (**A**) Pathway discovery performed via GSEA-based scoring (See **“Validation and replication of hnDAM signature”**) against hnDAM signature at varying z-score thresholds (z5 = 105 genes, z4 = 158, z3= 274, z2 = 521; **Table S51**) and other published signatures (**Table S53**). Curve plot of GSEA results for (**B**) hnDAM signature and (**C**) human microglia signature “Marshe_2025_GPNMB.hi”. Enrichment statistics available in **Table S52**.

